# C-LTMRs Regulate Thermosensation and Gate the Transition from Acute to Chronic Pain

**DOI:** 10.1101/2025.07.11.663895

**Authors:** Robert Guillaume, Karine Magalon, Aude Charron, Pascale Malapert, Chiara Salio, Andrew Saurin, Ana Reynders, Aziz Moqrich

## Abstract

C-low threshold mechanoreceptors (C-LTMRs) are traditionally associated with affective touch, yet emerging evidence suggests broader roles in sensory processing and pain modulation. We developed an intersectional genetic approach to selectively ablate C-LTMRs in adult mice by combining Nav1.8^IRES-FLPo^ and TH^CreER^ drivers with a conditional DTR reporter. This approach yields robust, tissue-specific deletion of C-LTMRs without off-target effects in non-sensory tissues. C-LTMR-ablated mice exhibit altered thermotaxis behavior, including a sharpened and spatially restricted preference for warmth, while maintaining largely intact responses to touch. Remarkably, following surgical or chemotherapeutic injury, these mice display persistent mechanical and cold hypersensitivity, implicating C-LTMRs in the resolution of pain. Transcriptomic profiling of dorsal root ganglia (DRG) and dorsal horn of the spinal cord (DHSC) revealed widespread transcriptional dysregulation in pathways related to extracellular matrix remodeling, vascular function and gliogenesis in naive mice. In C-LTMR-ablated mice, paclitaxel failed to induce pro-recovery transcriptional programs and instead promoted persistent neuroinflammatory signatures. These findings establish C-LTMRs as key modulators of pain recovery, acting through tissue-specific transcriptional programs that suppress inflammation and support sensory homeostasis.

## INTRODUCTION

Mechanical allodynia (pain elicited by normally innocuous mechanical stimuli) is a hallmark of several chronic pain conditions and remains a particularly challenging symptom to treat. A classical explanation for this phenomenon is provided by the gate control theory (GCT) of pain, first proposed by Melzack and Wall in 1965 ^1^. This model posits that tissue injury disrupts the inhibitory spinal “gate” circuits, allowing input from low-threshold Aβ mechanoreceptors to engage the pain pathway, thereby converting touch into pain. While this framework has guided decades of research, it may not fully capture the complexity of sensory modulation in chronic pain. Emerging evidence suggests that C-low-threshold mechanoreceptors (C-LTMRs), a class of unmyelinated sensory afferents traditionally associated with affective touch, may also play an active role in pathological pain states, potentially contributing to aberrant gating processes. C-LTMRs are specialized sensory neurons that respond preferentially to gentle stroking and innocuous cooling. Found in humans and other mammals, including cats and primates ^2,3^, their human counterparts, C-tactile afferents, are tuned to slow velocities (~3 cm/s) and skin temperature (~32°C), aligning with perceptions of pleasant touch^4,5^. These afferents terminate around hair follicles and project to lamina II of the dorsal horn, a key site for nociceptive and non-nociceptive integration ^6^. At the molecular level, C-LTMRs are marked by the expression of VGLUT3, tyrosine hydroxylase (TH), and TAFA4 ^7–13^, and are known to influence autonomic processes such as heart rate and to exert central effects through after-discharges and neuromodulation ^14^.

Importantly, C-LTMRs appear to have dual roles in pain modulation. Human studies show that their activation can produce both anti-nociceptive effects, such as reduced thermal or chemical pain ^15,16^, and pro-nociceptive responses, such as mechanical or cold allodynia in muscle pain and delayed-onset muscle soreness ^17–20^. Alterations in C-tactile function have also been observed in chronic pain conditions like migraine and fibromyalgia, suggesting a broader role in central sensitization ^21,22^.

Rodent studies have further explored the involvement of C-LTMRs in pain, but they suffer from important methodological limitations that complicate interpretation. For example, initial findings using VGLUT3-KO mice suggested that C-LTMRs contribute to mechanical hypersensitivity ^12^, but follow-up work showed this phenotype was due to transient VGLUT3 expression in spinal neurons, not in C-LTMRs themselves ^23^. Similarly, conditional inactivation of Cav3.2 in Nav1.8-lineage neurons, intended to target Cav3.2 function in C-LTMRs, reduced pain responses ^24^, but the broader expression of this marker in other C-fiber subsets undermines the specificity of these conclusions. Optogenetic studies also face similar challenges. Noble et al. (2022) showed that activation of TH-lineage neurons in injured mice induced aversive behaviors, implicating a pronociceptive role for C-LTMRs ^25^. However, TH is also expressed in sympathetic fibers, leaving open the possibility that the observed effects were not specific to C-LTMRs. In the context of chemotherapy-induced pain, optogenetic stimulation of VGLUT3-lineage cells produced nocifensive behaviors in oxaliplatin-treated mice ^26^, but again, lineage tracing does not guarantee functional specificity to C-LTMRs.

On the other hand, several studies using TAFA4 and Bhlha9 KO mice as well as GINIP-DTR mice have proposed anti-nociceptive roles for C-LTMRs ^7,27–29^, suggesting these neurons may also contribute to the resolution of pain. Yet, these findings rely on global gene deletions, dual ablation of C-LTMRs and MRGPRD^+^ mechanonociceptors or exogenous protein application, all of which fail to isolate the endogenous and cell-specific contributions of C-LTMRs.

Taken together, while experimental data from both human and animal studies suggest that C-LTMRs may exert bidirectional effects on pain, current approaches, including global and conditional knock-out models, lack the cellular and temporal specificity necessary to definitively link C-LTMRs to specific pain phenotypes. These limitations highlight the need for more precise, lineage-restricted, and temporally controlled strategies to clarify the true functional role of C-LTMRs in pain modulation. In this study, we developed a novel, inducible genetic model to selectively ablate C-LTMRs in adult mice. By crossing *Nav1.8^IRES-FLPo^*;*TH^CreER^*mice with a dual-recombinase reporter line expressing diphtheria toxin receptor (DTR), we achieved robust, tissue-specific ablation of C-LTMRs. This inducible genetic strategy circumvents the off-target effects and developmental compensations commonly associated with traditional knockout models. Using this approach, we demonstrate that partial ablation of C-LTMRs in adult mice does not substantially impair touch behaviors but markedly alters thermotaxis preferences and significantly promotes the transition from acute to chronic pain following both surgical and chemotherapeutic injury. Transcriptomic analyses of DRG and spinal cord tissues confirmed the selective loss of C-LTMR-specific markers and revealed broad dysregulation of genes involved in synaptic transmission, thermogenesis, immune signaling, and detoxification pathways. These findings establish C-LTMRs as key modulators of thermal preference and critical contributors to the recovery processes after injury.

## RESULTS

### Tissue specific and inducible ablation of C-LTMRs

Previous studies from our laboratory have suggested that, beyond their well-established role in mediating pleasant touch, C-LTMRs may also modulate pain following tissue injury ^7,28^. To directly test this hypothesis, we developed a genetic strategy enabling the selective and inducible ablation of C-LTMRs in adult mice. We crossed Na_v_1.8^Ires−FLPo^;TH^CreER^ mice with a reporter line expressing the simian diphtheria toxin receptor (DTR) under the control of a pan-neuronal *Tau* promoter flanked by loxP and FRT stop cassettes (Tau^loxP-STOP-loxP-FRT-STOP-FRT-DTR^) (Fig. 1A). This breeding strategy yielded two genotypes: experimental Na_v_1.8^Ires−FLPo^;TH^CreER^;Tau^DTR^ (hereafter referred to as *C-LTMRs-DTR* mice) and control Na_v_1.8^Ires−FLPo^;Tau^DTR^ (hereafter referred to as control mice).

**Figure 1.**
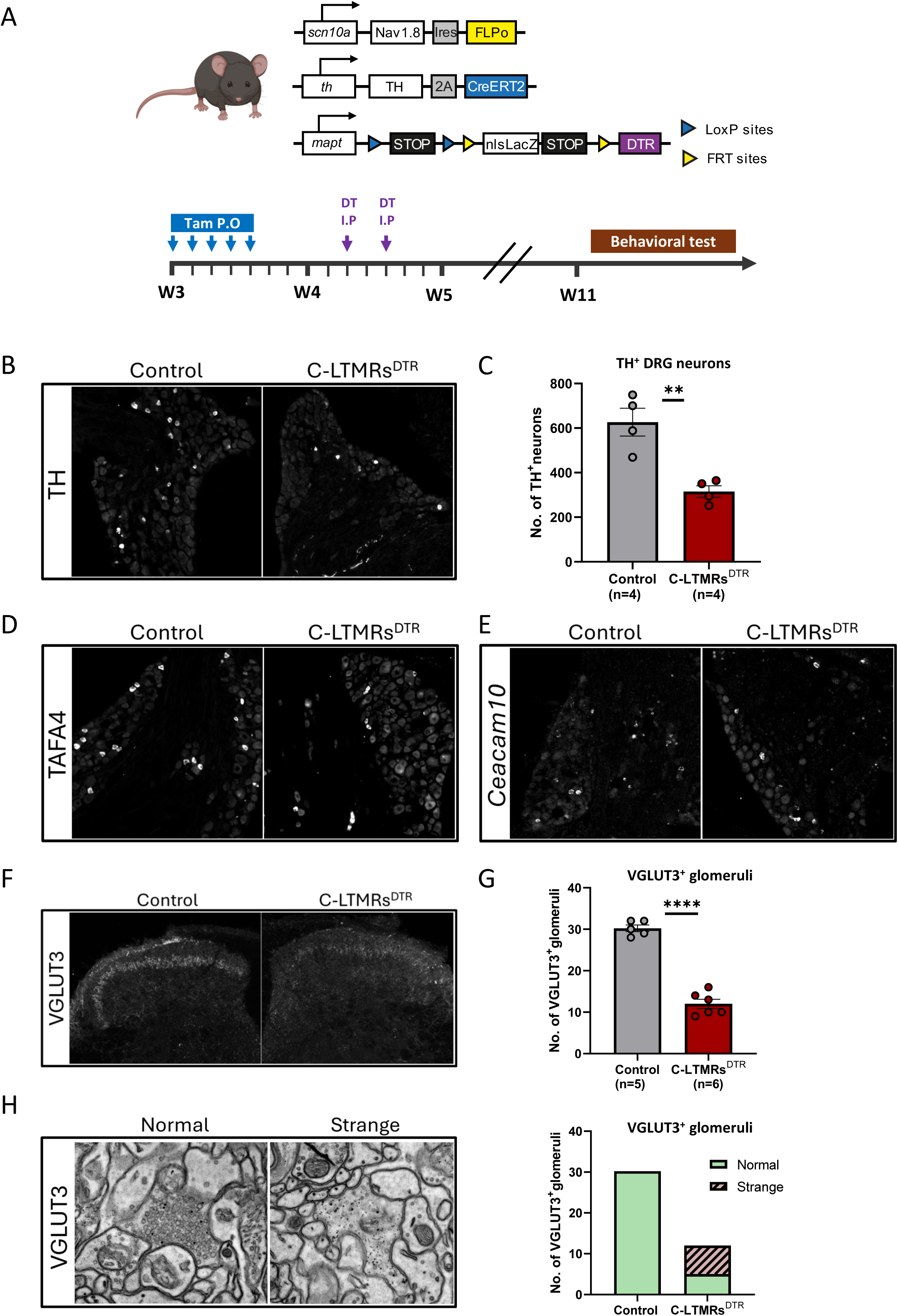
Selective and inducible ablation of C-LTMRs in adult mice. **(A)** Schematic of the intersectional genetic strategy used to selectively ablate C-LTMRs. *Nav1.8^IRES-FLPo^* drives expression of FLPo in sensory neurons; *TH^CreER^* enables tamoxifen-inducible Cre expression in TH^+^ neurons. The *Tau-^loxP-STOP-loxP-FRT-STOP-FRT-DTR^* allele allows DTR expression only in neurons where both Cre and FLPo are active, permitting C-LTMR-specific ablation upon diphtheria toxin (DT) administration. **(B)** Representative images of lumbar (L3) DRG sections stained TH in control and *C-LTMRs-DTR* mice. **(C)** Quantification of the total number of lumbar (L3) TH^+^ DRG neurons in control (grey) and *C-LTMRs-DTR (red)* mice (*n* = 4 per group). C-LTMR ablation results in a ~50% reduction in TH^+^ neurons. Data are presented as the mean ± SEM (unpaired t-test *\*\*P < 0.01).* Dots represent individual animals. **(D-E)** Expression levels of C-LTMRs-enriched markers (TAFA4 and *Ceacam10*) are significantly decreased in *C-LTMRs-DTR* mice compared to control mice. **(F)** Immunostaining for VGLUT3 in the spinal cord dorsal horn. *C-LTMRs-DTR* mice show a dramatic loss of VGLUT3 signal in lamina II inner compared to controls. **(G)** Quantification of VGLUT3^+^ glomeruli in the DHSC, in control (gray, n=5) and *C-LTMRs-DTR* mice (red, n=6). Data are presented as the mean ± SEM (unpaired t-test *\*\*\*\*P <0.0001).* Dots represent individual animals. **(H)** Ultrastructure of C-LTMRs terminals immuno-gold labelled for VGLUT3. Strange terminals, only observed in *C-LTMRs-DTR* mice, exhibit a drastic decrease in the number of synaptic vesicles compared to normal ones.

To induce targeted neuronal ablation, we administered tamoxifen (oral gavage, once daily for five consecutive days) to 3-week-old mice, followed one week later by two intraperitoneal injections of diphtheria toxin (DT) spaced two days apart. All subsequent analyses were conducted at 11 weeks of age (Fig. 1A).

To assess the efficacy and specificity of C-LTMRs ablation, we quantified the number of tyrosine hydroxylase-positive (TH^+^) neurons in lumbar (L3) dorsal root ganglia (DRG) (Fig. 1B and C). Compared to controls, *C-LTMRs-DTR* mice exhibited an approximately 50% reduction in TH+ neuron counts (Fig. 1C) (control: 625,5±62,38; *C-LTMRs-DTR*: 315±25.88; *n* = 4). Consistently, expression of C-LTMRs-enriched markers such as TAFA4 and *Ceacam10* (Delfini et al., 2013; Reynders et al., 2015) was markedly reduced in *C-LTMRs-DTR* mice (Fig. 1D and E). In contrast, markers excluded from C-LTMRs, including CGRP, P2X3, IB4, TrkC, and NF200 remained unchanged (Fig. S1A and B), supporting the selectivity of our ablation approach.

At the spinal level, *C-LTMRs-DTR* mice showed a pronounced decrease of VGLUT3 immunoreactivity specifically in lamina II inner of the dorsal horn (Fig. 1F). This was corroborated by electron microscopy, which revealed a dramatic reduction in VGLUT3^+^ glomeruli in *C-LTMRs-DTR* mice compared to controls (Fig. 1G). This is consistent with the large decrease of the number of non-labelled glomeruli observed in *C-LTMRs-DTR* mice (Fig. S1C). Note that a mild but significant decrease in IB4^+^ glomeruli was also observed (Fig S1D). Additionally, the analysis revealed the presence of some VGLUT3^+^ glomeruli with atypical morphology, referred to as “strange” glomeruli, which exhibited a markedly reduced density of synaptic vesicles compared to typical (“normal”) glomeruli (Fig. 1H). These “strange” glomeruli were only observed in *C-LTMRs-DTR* mice (Fig. 1H) suggesting a functional deficit in some of the 50% remaining VGLUT3^+^ glomeruli.

To rule out off-target effects in non-sensory tissues, we crossed *Nav1.8^Ires-FLPo^* mice with the *RC::FL-hM3Dq* reporter line ^30^, allowing visualization of *Nav1.8*-lineage via EGFP expression (Fig. S2A). EGFP signal was absent from known TH^+^ regions such as Substantia Negra pars compacta (SNpc) in the brain (Fig. S2B), jugular/nodose ganglion (JNG) in which a faint colocalization signal is occasionally detectable (Fig. S2C), and celiac ganglia (Fig. S2D). These data confirm that *Nav1.8*-driven FLPo activity, and thus C-LTMR ablation, is restricted to primary sensory neurons. Together, these results demonstrate that our genetic model enables robust, inducible, and tissue-specific ablation of C-LTMRs in adult mice. This approach provides a powerful tool for dissecting the functional contributions of C-LTMRs to somatosensation and pain modulation.

### Partial ablation of C-LTMRs alters thermotaxis behavior without significantly affecting touch responses

We first sought to assess the impact of partial ablation of C-LTMRs on general behavior. *C-LTMRs-DTR* mice appeared normal in terms of open field (Figure S3A), and rotarod (Figure S3B) profiles, indicating that a 50% reduction in C-LTMRs does not result in detectable alterations in motor activity or anxiety-like behavior.

While C-LTMRs are classically associated with affective touch, several studies have suggested they may also play a role in temperature perception ^6,24,27^. To directly investigate their contribution to thermosensation, we assessed thermotaxis behavior of *C-LTMRs-DTR* mice using a temperature gradient paradigm.

Over a 90-minute session, *C-LTMRs-DTR* mice of both sexes exhibited a significantly sharper and more spatially restricted preference for warmer temperatures compared to their control littermates (Fig. 2A and 2B). When we analyzed behavior in the first 30-minute interval, all groups explored the arena in a similar manner, indicating normal exploratory drive (Fig. 2C and D). By the second interval, both control and *C-LTMRs-DTR* males began favoring warmer zones, but this preference was significantly more pronounced in ablated mice and further intensified during the last 30 minutes (Fig. 2C). A similar, yet more striking, effect was observed in females: while control females continued to explore a broader range of temperatures, *C-LTMRs-DTR* females rapidly developed and maintained a strong preference for warmer areas, closely mirroring the male ablation phenotype (Fig. 2D).

**Figure 2.**
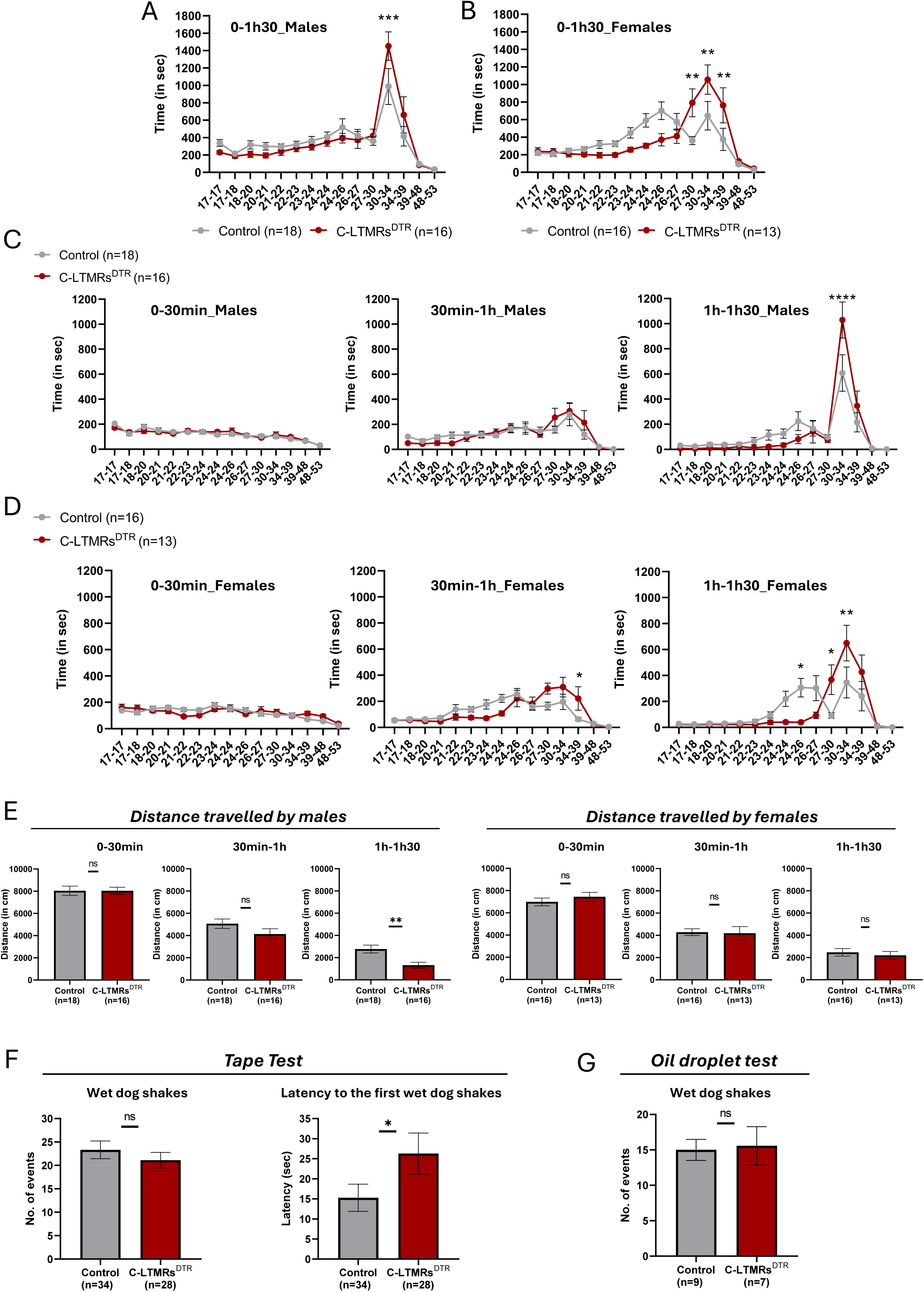
Partial ablation of C-LTMRs sharpens thermotactic behavior without major disruption of gentle touch responses. **(A-B)** Graphical representations of the time spent by (A) males and (B) females control (gray line) and *C-LTMRs-DTR* mice (red line) across a 90-minute temperature gradient assay. Ablated mice of both sexes (n=16 for males and n=13 for females) show a markedly narrower preference for the warm zone compared to controls (n=18 for males and n=16 for females). Data are presented as mean ± SEM (two-way repeated measures ANOVA followed by Bonferroni test) **(C)** Temporal analysis across three 30-minute intervals shows that both male groups initially explore the full arena, but *C-LTMRs-DTR* males progressively restrict their movements to the warmer zone, showing a significantly stronger thermal preference than controls by the final phase. Data are presented as mean ± SEM (two-way repeated measures ANOVA followed by Bonferroni test). **(D)** Female *C-LTMRs-DTR* mice show an even earlier and more pronounced spatial restriction to the warm zone compared to control females, who maintain broader temperature exploration. Data are presented as mean ± SEM (two-way repeated measures ANOVA followed by Bonferroni test). **(E)** Total distance traveled in each phase reveals no difference between genotypes during initial exploration. By the final 30 minutes, *C-LTMRs-DTR* males travel significantly less than controls, consistent with increased spatial fixation. No significant differences were observed in females. Data are presented as mean ± SEM (unpaired Mann-Whitney test or t-test, ns, ** P < 0.01). **(F-G)** Behavioral responses to gentle touch were evaluated using the tape removal test and the oil drop assay. No significant differences were observed between groups in the number of wet dog shakes response. However, *C-LTMRs-DTR* mice exhibited a slight but consistent delay in the onset of the “wet-dog shake” response in the tape test, suggesting altered C-LTMR signaling dynamics (Zhang et al., *Science*, 2024). Data are presented as mean ± SEM (unpaired Mann-Whitney test or t-test, ns, * P < 0.05).

To ensure this thermotaxis phenotype was not confounded by motor deficits, we quantified total distance traveled across each phase. All groups covered comparable distances during the first 30 minutes (control males: 8041±412,1 cm, C-LTMRs^DTR^ males: 8039±318,3cm, control females: 6984±343cm, C-LTMRs^DTR^ females: 7443±403,1cm), confirming intact locomotor function (Fig. 2E). During the second and third intervals, movement decreased in all groups as temperature preferences consolidated (Fig. 2E). Notably, *C-LTMRs-DTR* males traveled significantly less than controls in the final phase (1325 ± 258.6 cm vs. 2774±354,3cm), consistent with significantly higher sharp and net preference for the warm zone of the arena (Fig. 2E). No significant difference was observed between female groups, likely due to broader zone occupancy across genotypes (Fig. 2E).

Next, we sought to evaluate whether the partial ablation of C-LTMRs also affects touch responses. To do so, we assessed behavioral outcomes using the tape removal ^31^and the oil drop assay ^32^. In both paradigms, *C-LTMRs-DTR* mice performed similarly (in terms of number of wet dog shakes) to controls, indicating that basic touch perception remains largely intact under our C-LTMRs’ ablation conditions (Fig 2F and G). However, in the tape test, we observed a mild but consistent delay in the onset of the characteristic “wet-dog shake” response in *C-LTMRs-DTR* mice (Figure 2F); a behavior recently linked to C-LTMR-mediated activation of the spinoparabrachial pathway ^32^.

Together, these findings demonstrate a novel role for C-LTMRs in shaping thermotaxis behavior and spatial temperature discrimination, while suggesting that the baseline detection of touch may either require the full complement of C-LTMRs or depend on subpopulations that remain intact in our ablation set up.

### Partial ablation of C-LTMRs facilitates the transition to chronic pain following injury

In earlier work, we identified GINIP (Gα-inhibitory interacting protein) as a marker of two distinct populations of dorsal root ganglion (DRG) neurons: MRGPRD-expressing mechanonociceptors and TAFA4-expressing C-LTMRs ^33^. Using a genetic ablation approach targeting both populations simultaneously, we later showed that *GINIP-DTR* mice displayed normal onset and resolution of mechanical hypersensitivity in both the Complete Freund’s Adjuvant (CFA) inflammatory model and the chronic constriction injury (CCI) model of neuropathic pain ^28^. We hypothesized that this lack of phenotype could be due to compensatory effects between the two ablated populations: specifically, that ablation of MRGPRD+ neurons alone would be analgesic ^34^, while ablation of C-LTMRs alone would exaggerate injury-induced pain. Thus, the concurrent deletion of both populations may have masked the individual contributions of each.

To directly test the specific role of C-LTMRs in injury-induced pain, we examined *C-LTMRs-DTR* and control mice in two complementary models: the paw incision surgery as a model of acute post-operative pain ^35^ and paclitaxel-induced chemotherapy-induced peripheral neuropathy (CIPN), a model of chronic neuropathic pain ^36^. Mechanical sensitivity was assessed at baseline and multiple time points post-injury using the Von Frey up-and-down method.

At baseline, both male and female *C-LTMRs-DTR* mice exhibited a modest but significant reduction in mechanical thresholds compared to controls (Fig. 3A; *n* = 43 and 44, respectively and S4A). To account for this difference, all subsequent data were normalized to individual baselines and expressed as percentage changes.

**Figure 3.**
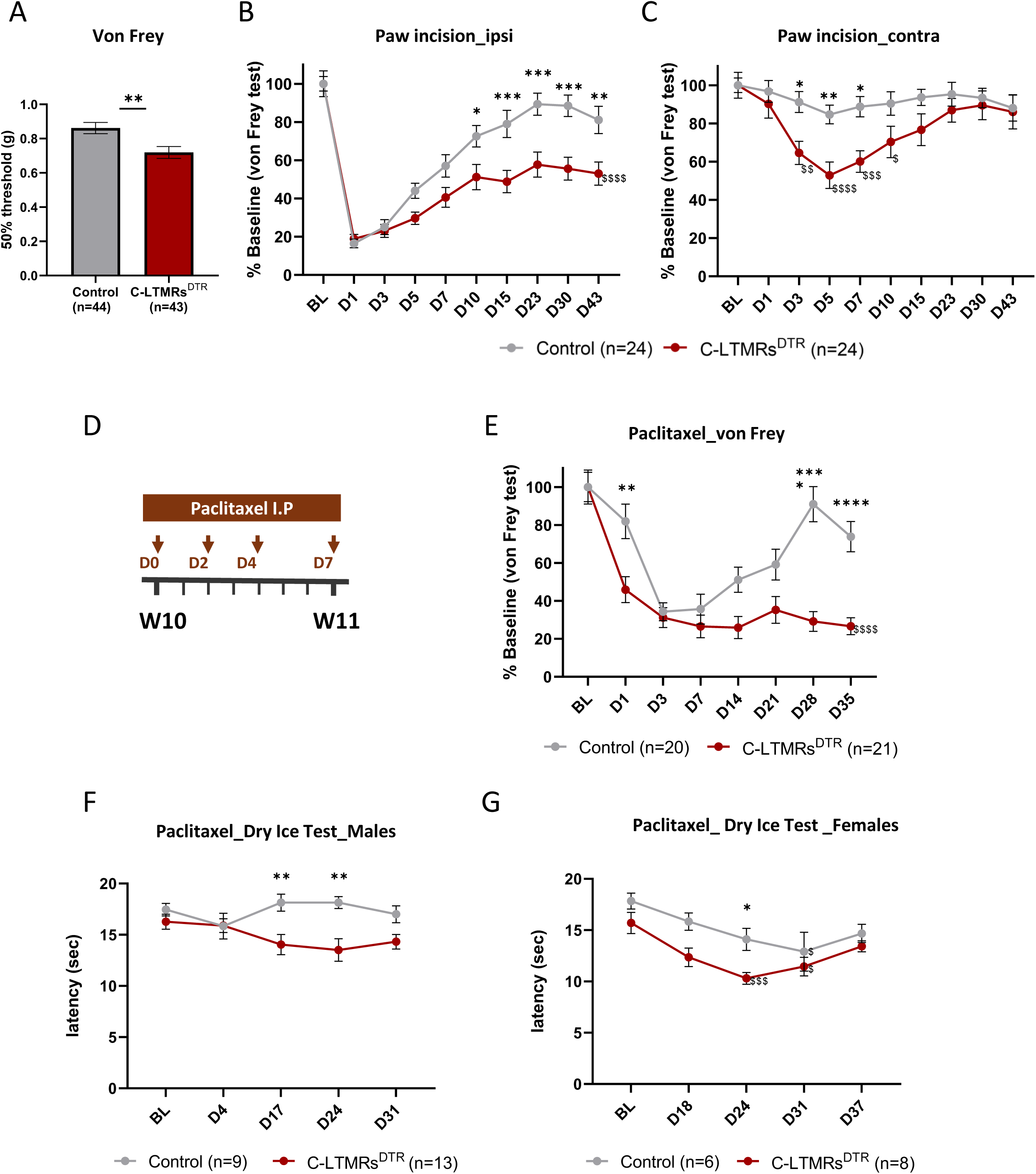
Partial ablation of C-LTMRs prolongs injury-induced mechanical hypersensitivity and promote cold allodynia. **(A)** Baseline mechanical thresholds measured by Von Frey testing show a modest but significant reduction in *C-LTMRs-DTR* mice compared to controls (*n* = 43 and 44, respectively) (unpaired t-test ** P < 0.01). **(B)** In the paw incision model, both groups develop acute mechanical hypersensitivity at day 1 post-surgery. However, *C-LTMRs-DTR* mice fail to recover and display persistent hypersensitivity lasting up to 43 days (*n* = 24 per group) (two-way repeated measures ANOVA followed by Bonferroni test). **(C)** Repetitive mechanical stimulation of the contralateral paw reveals a significant reduction in threshold in *C-LTMRs-DTR* but not control mice, indicating generalized sensitization (two-way repeated measures ANOVA followed by Bonferroni test). **(D)** Schematic of the paclitaxel-induced CIPN model. **(E)** *C-LTMRs-DTR* mice show an earlier and more severe mechanical hypersensitivity that persists beyond the treatment period (control: *n* = 20, C*-LTMRs-DTR: 21*) (two-way repeated measures ANOVA followed by Bonferroni test). **(F-G)** Cold allodynia develops selectively in *C-LTMRs-DTR* mice after paclitaxel administration, with significant threshold reduction starting around day 17 for males, and day 24 for females. All data are presented as mean ± SEM, *P < 0.05, **P < 0.01, ***P < 0.001 ***P < 0.0001, idem for $ indicating statistical differences in *C-LTMRs-DTR* mice (or control mice only Fig. 3G) between D0 and the corresponding time point.

In the paw incision model, both *C-LTMRs-DTR* and control mice developed robust mechanical hypersensitivity in the ipsilateral hindpaw by day 1 post-surgery (Fig. 3B; *n* = 24 per group). While hypersensitivity gradually resolved in control animals, *C-LTMRs-DTR* mice displayed a persistent pain phenotype that lasted up to 43 days, indicating impaired recovery (Fig. 3B). Moreover, repetitive stimulation of the contralateral (uninjured) paw revealed a significant reduction in mechanical threshold in *C-LTMRs-DTR* mice, but not in controls (Fig. 3C), suggesting an enhanced wind-up phenomenon or a generalized sensitization phenotype. No statistical differences were observed between males and females of both genotypes neither for the ipsilateral side nor the contralateral one. (Fig. S4B and C).

To determine whether this effect extended to neuropathic pain, we employed the paclitaxel CIPN model (Fig. 3D). In both groups, paclitaxel administration induced mechanical hypersensitivity (Fig. 3E). However, *C-LTMRs-DTR* mice developed a significantly earlier onset and more severe mechanical hypersensitivity than controls, which persisted well beyond the treatment window, showing no signs of resolution (Fig. 3E). No statistical differences were observed between males and females of both genotypes (Fig. S4D).

Given that paclitaxel is also known to induce cold allodynia, we assessed cold sensitivity using the dry ice test ^37^. First at baseline level, no differences were observed between both genotypes and nor between sexes in each genotype (Fig. S4E). Repetitive paclitaxel injections led to a significant increase in cold sensitivity in *C-LTMRs-DTR* mice compared to controls starting at day 17 post-treatment (Fig. 3F and G).

Taken together, these findings reveal a previously unappreciated role for C-LTMRs in the resolution of injury-induced pain. Their partial ablation delays recovery and promotes the transition from acute to chronic pain, suggesting that C-LTMRs act as modulators of protective sensory signaling during the recovery process.

### Partial ablation of C-LTMRs deregulate transcriptional programs associated with recovery from chemotherapy-induced pain

To investigate the molecular mechanisms by which C-LTMR ablation impairs thermotaxis and promotes mechanical hypersensitivity upon repeated stimulation, we performed bulk RNA sequencing on naïve adult L3–L5 dorsal root ganglia (DRG) and dorsal horn of the spinal cord (DHSC) from control and *C-LTMRs-DTR* mice (Fig. 4 and 5).

**Figure 4.**
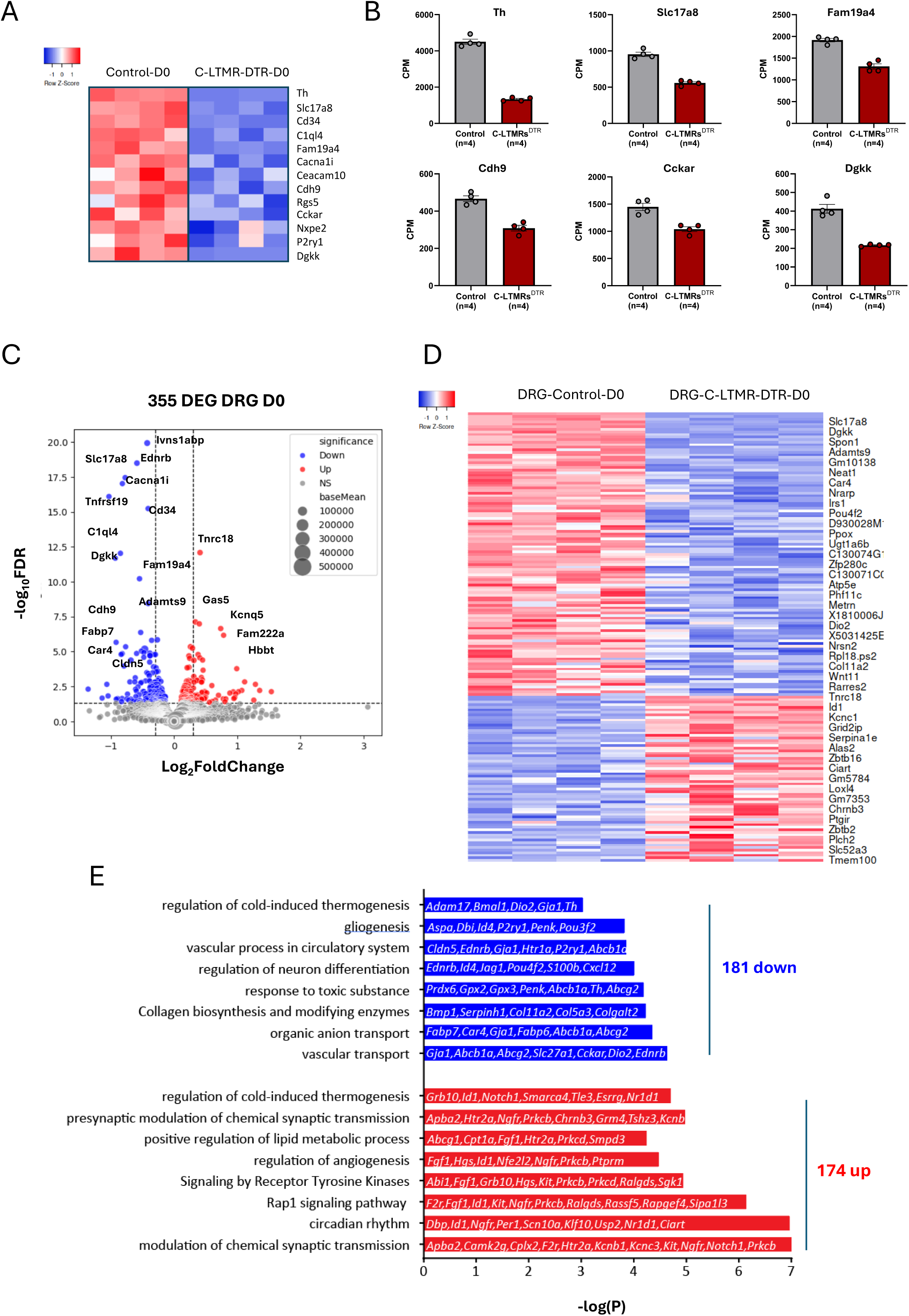
Partial ablation of C-LTMRs modifies gene expression in the lumbar DRGs in naive mice. A) Heatmap representing the expression of indicated C-LTMRs-specific or enriched genes in Control and *C-LTMRs-DTR* naïve mice. B) Counts Per Million (CPM) obtained after RNA-Seq for the same sets of C-LTMR-specific or enriched genes. C) Volacno plot representing up- and down-regulated DEG in the L3-L5 DRG of naive *C-LTMRs-DTR* compared to Control mice (FDR5, n=4 samples per genotype). For a matter of representation, *Th* was excluded from the plot. D) Heatmap representing the expression of identified DEG in Control and *C-LTMRs-DTR* mice (n=4 each). E) Up (red) and down (blue)-regulated biological processes and pathways in *C-LTMRs-DTR* lumbar DRGs determined with Metascape software. Non-exhaustive examples of corresponding transcripts are illustrated for each indicated processes.

**Figure 5.**
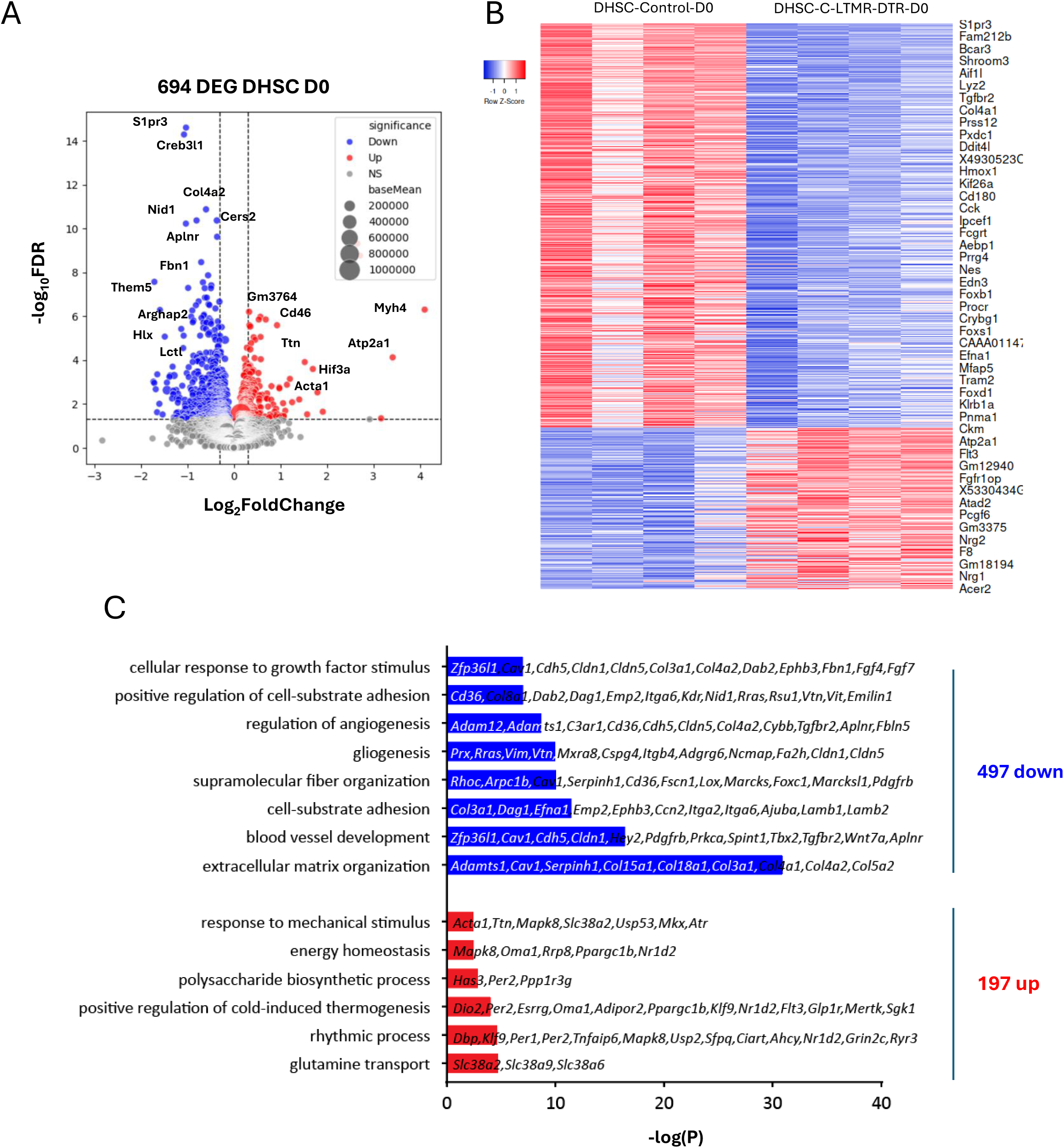
Partial ablation of C-LTMRs modifies gene expression in the lumbar DHSC in naive mice. A) Volacno plot representing up- and down-regulated DEG in the lumbar DHSC of naive *C-LTMRs-DTR* mice compared to Controls (FDR5, Log2FoldChange > 0.3, n=4 samples per genotype). B) Heatmap representing the expression of identified DEG in naïve control and *C-LTMRs-DTR* mice (n=4 each). E) Up (red) and down (blue)-regulated biological processes and pathways in *C-LTMRs-DTR* lumbar DHSC determined with Metascape software. Non-exhaustive examples of corresponding transcripts are illustrated for each indicated processes.

Differential gene expression analysis confirmed successful ablation of C-LTMRs, evidenced by a marked reduction in transcripts highly enriched in this population, including *Th*, *Fam19a4* (TAFA4), *Slc17a8* (VGLUT3), *Cdh9, Cckar, Dgkk* and many others (Fig. 4A-B and S5A).

To identify molecular signatures associated with altered thermotaxis and mechanical hypersensitivity in the absence of injury, we compared gene expression profiles between control and *C-LTMRs-DTR* mice. In DRGs, we identified 355 differentially expressed genes (DEGs) at a 5% false discovery rate (FDR), with 174 genes upregulated and 181 downregulated in *C-LTMRs-DTR* mice (Fig. 4C-E). Gene ontology (GO) analysis of upregulated DEGs revealed enrichment in biological processes related to synaptic transmission, circadian rhythm, Rap1 signaling, and cold-induced thermogenesis (Fig. 4E). In contrast, downregulated genes were associated with vascular and organic anion transport, collagen biosynthesis, responses to toxic substances, and neuronal/glial differentiation (Fig. 4E).

In the DHSC, transcriptomic analysis revealed 694 DEGs (197 upregulated, 497 downregulated; FDR 5%, |log2FC| ≥ 0.3), indicating that C-LTMR ablation substantially alters the spinal transcriptional landscape under naïve conditions (Fig. 5A-C). Similar to DRGs, upregulated genes were associated with thermogenesis and mechanical stimulus responses (Fig. 5C). Additionally, genes linked to glutamine metabolism, energy homeostasis, and polysaccharide biosynthesis were upregulated (Fig. 5C). Strikingly, downregulated genes were enriched in processes related to extracellular matrix (ECM) organization, tissue morphogenesis, blood vessel development, and gliogenesis (Fig. 5C).

These findings suggest that in naïve mice, the absence of C-LTMRs not only alters neuronal signalling but also disrupts broader metabolic, vascular, and glial homeostasis in both DRG and DHSC tissues, potentially contributing to altered sensory behavior.

To further explore how C-LTMRs influence the development of chronic pain, we performed bulk RNA sequencing on ipsilateral L3–L5 DRGs and lumbar DHSC at baseline (Day 0) and 40 days after paclitaxel treatment (Day 40) in control and *C-LTMRs-DTR* mice

As a control, we first confirmed the effective ablation of C-LTMRs, in *C-LTMRs-DTR* mice subjected to paclitaxel treatment (Fig. S5B). Consistent with previous observations, we detected a substantial reduction in the expression of genes highly enriched in C-LTMRs, including *Th*, *Slc17a8* (VGLUT3), *Fam19a4* (TAFA4), *Cacna1i* (Cav3.3) and *Ceacam10* (Fig. S5B). In DRGs from control mice, 178 DEGs were identified post-paclitaxel (48 upregulated, 130 downregulated; FDR 5%, |log2FC| ≥ 0.3), compared to 107 DEGs in *C-LTMRs-DTR* mice (67 upregulated, 40 downregulated), with minimal overlap between genotypes (Fig.S6 A-D). In line with previous studies ^38,39^, paclitaxel treatment in control mice upregulated genes related to circadian regulation, thermogenesis, and hypoxia response, while downregulating genes involved in ECM remodelling, skeletal and skin development, and cytokine production (Fig. S6B). These transcriptional changes are consistent with adaptive DRG responses that likely support recovery from mechanical and cold hypersensitivity.

In contrast, DRGs of *C-LTMRs-DTR* mice showed distinct transcriptional responses to paclitaxel. Although ECM-related processes were also suppressed, we observed a unique upregulation of genes involved in macrophage activation and innate immune responses, suggesting that C-LTMRs play a role in restraining neuroinflammatory pathways (Fig. S6D). Transcriptomic analysis of DHSC samples revealed even more pronounced differences. In control mice, 479 DEGs (399 downregulated, 80 upregulated, FDR 5, |log2FC| ≥ 0.3) were detected at Day 40 post-paclitaxel (Fig. S7A-B). Strikingly, *C-LTMRs-DTR* mice exhibited 1,059 DEGs (836 downregulated, 223 upregulated,FDR 5, |log2FC| ≥ 0.3), indicating that the absence of C-LTMRs profoundly alters the spinal response to chemotherapy (Fig. S7A-B).

Among these, only 116 DEGs were shared between genotypes, suggesting a core set of genes consistently affected by paclitaxel (Fig. S7 A-C). Further comparisons highlighted major transcriptional programs disrupted specifically in *C-LTMRs-DTR* mice. Notably, 122 genes were selectively upregulated, associated with neutrophil activation, cell proliferation, and secretory processes (Fig S7B, S8A-B, 6A-B). Conversely, 654 genes were strongly downregulated, with enrichment for processes involved in gene regulation and calcium ion transport (Fig S7A, S8C-D, 6A-B). Additionally, 292 genes that were significantly downregulated in control mice post-paclitaxel were not similarly regulated in *C-LTMRs-DTR* mice (Fig. S9A-B). These genes were predominantly associated with ECM and vascular remodelling and had already shown reduced expression in naïve *C-LTMRs-DTR* mice (Fig. 6A and C). This suggests that the lack of C-LTMRs impairs the ability to implement reparative transcriptional responses likely necessary for recovery.

**Figure 6.**
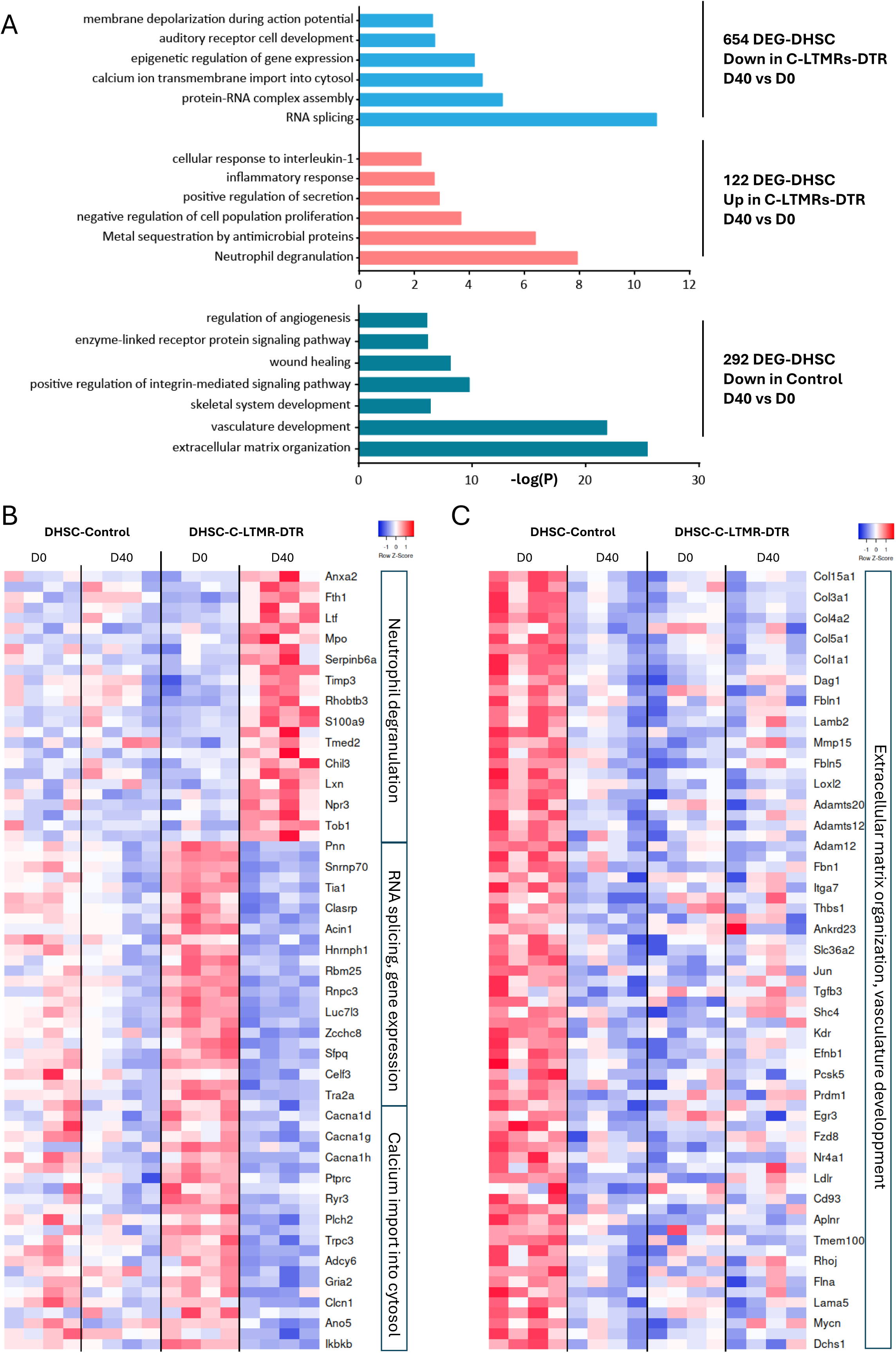
Partial ablation of C-LTMRs disrupts DHSC transcriptional response to paclitaxel. A) Down-(light blue, upper panel) and up-(light red, middle panel) regulated biological processes and pathways in *C-LTMRs-DTR* lumbar DHSC at day 40 (D40) after paclitaxel as compared to untreated *C-LTMRs-DTR* DHSC (D0). Down-regulated (dark turquoise, lower panel) biological processes and pathways in Control lumbar DHSC at D40 after paclitaxel as compared to untreated Control DHSC (D0). B) Heatmap representing the expression levels of genes involved in *Neutrophil degranulation*, *RNA-Splicing and gene expression*, *Calcium import in the cytosol* in Control and *C-LTMRs-DTR* untreated (D0) and at D40 after paclitaxel administration (D40). These genes are part of transcriptional changes occurring specifically in *C-LTMRs-DTR* DHSC in response to paclitaxel (see Fig.S8). C) Heatmap representing the expression levels of genes involved in Extracellular matrix organization and vasculature development in Control and *C-LTMRs-DTR* untreated (D0) and at D40 after paclitaxel administration (D40). These genes are part of the transcriptional response to paclitaxel implemented in Controls but not in *C-LTMRs-DTR* DHSC (see Fig.S9).

Together, these data reveal that in the absence of C-LTMRs, both DRG and DHSC tissues enter a transcriptionally altered state affecting key metabolic, detoxification, and vascular processes. These alterations compromise the tissue’s ability to mount adaptive responses to paclitaxel, promoting prolonged hypersensitivity and a failure to engage pro-recovery pathways. In sum, C-LTMRs are essential for orchestrating transcriptional programs that foster resolution of pain through ECM remodelling and by suppressing neuroinflammatory signalling.

## DISCUSSION

C-LTMRs have been classically associated with affective touch and social affiliative behaviors. In this study, we uncover a broader, previously underappreciated role for C-LTMRs in maintaining sensory homeostasis and regulating the transition from acute to chronic pain. By employing a tamoxifen-and diphtheria toxin-inducible genetic strategy, we achieved selective and tissue-specific ablation of C-LTMRs in adult mice. This model enabled us to dissect their contributions beyond touch, revealing key functions in thermosensation, injury resolution, and pain modulation.

Although historically linked to pleasant touch, accumulating evidence suggests that C-LTMRs also contribute to innocuous thermal perception. Prior in vitro and ex vivo work has shown that C-LTMRs respond to gradual temperature changes, particularly across broad cooling and warming ranges, without signaling discrete thresholds ^6,24^. Our behavioral findings support this thermosensory function in vivo.

In a thermal gradient assay, mice with partial C-LTMR ablation exhibited a significantly sharper and more spatially restricted preference for warmer temperatures compared to controls. This phenotype was consistent across sexes, emerged gradually, and was not attributable to deficits in locomotion or broad sensory impairment, as exploratory behavior and responses to noxious temperatures were preserved. These findings suggest that C-LTMRs contribute to the detection of relative temperature changes and help guide adaptive thermotaxis behavior. Unlike classical thermoreceptors, which rely on TRP channel-mediated detection of absolute thermal thresholds, C-LTMRs may provide a continuous, graded input that enhances subtle temperature discrimination. In humans, the pleasantness of touch is known to peak at skin temperature (~32°C), where CT afferents (the human equivalents of C-LTMRs) exhibit maximal firing ^4^. This convergence of tactile and thermal tuning suggests that C-LTMRs encode not just mechanical features of touch, but also its thermal context.

Despite this clear thermosensory phenotype, baseline tactile responses in *C-LTMRs-DTR* mice were largely preserved. In assays such as the tape removal test and oil drop assay, ablated animals performed comparably to controls. A minor but consistent delay in the wet-dog shake reflex was observed, a behavior recently linked to C-LTMR-mediated engagement of the spinoparabrachial pathway ^32^. These findings suggest that partial ablation either spares a sufficient number of C-LTMRs to support basic tactile perception or that redundant pathways involving other low-threshold mechanoreceptors compensate for the loss. Supporting the former hypothesis, single-cell RNA sequencing has identified two subsets of C-LTMRs distinguished by differing levels of tyrosine hydroxylase (TH) expression ^8^. Given that the *Th* locus drives Cre recombinase expression in this model, it is plausible that the spared population contributing to residual tactile behaviors, including tape removal and the wet-dog shake, is the subset expressing lower levels of TH.

A more striking phenotype emerged under conditions of repeated mechanical stimulation. *C-LTMRs-DTR* mice exhibited increased mechanical hypersensitivity in both acute and chronic pain models, including exaggerated responses to repetitive stimulation of uninjured tissue; a pattern resembling wind-up ^40^. These data suggest that C-LTMRs are involved in suppressing this amplification process under normal conditions. This interpretation is supported by human psychophysical studies demonstrating that CT-targeted touch reduces wind-up pain and modulates cortical responses associated with central sensitization. For instance, Fidanza et al. (2021) showed that gentle stroking of CT-innervated skin inhibits wind-up, while similar stimuli to glabrous skin do not ^41^. Wakui et al. (2025) further demonstrated that CT activation uniquely shapes cortical responses to repetitive mechanical input ^42^. Similarly, Taneja et al. (2021) found that continuous CT-optimal stimulation alleviates hyperalgesia on remote body regions, but only when delivered with CT-specific parameters ^43^. Together, these findings support a conserved role for C-LTMRs in damping excitatory spinal signaling and modulating central pain states.

The most profound effect of C-LTMR ablation was observed following injury. In both paw incision and Paclitaxel-induced neuropathy models, *C-LTMRs-DTR* mice failed to recover from mechanical hypersensitivity and instead developed persistent pain. This occurred despite intact Aβ fibers and other low-threshold mechanoreceptors, underscoring a protective role for C-LTMRs in pain resolution and strongly implicating this enigmatic population in the gate control theory of pain ^1^. While this theory has long provided a foundational framework in pain research, the specific afferent subtypes mediating gating mechanisms have remained elusive. Our findings provide direct evidence that C-LTMRs contribute to this gating function by modulating pain resolution after injury. Rather than broadly inhibiting nociceptive input, C-LTMRs appear to selectively constrain maladaptive plasticity during the transition from acute to chronic pain. Their loss unmasks latent sensitization, thereby promoting the development of prolonged pain states.

How, mechanistically speaking, C-LTMRs contribute to the development of prolonged pain states? Transcriptomic analyses revealed that C-LTMR ablation leads to substantial dysregulation of gene expression in both DRG and DHSC, even under naïve conditions. Key findings included downregulation of genes involved in extracellular matrix organization and vascular remodeling, likely impairing the structural and metabolic support required for sensory circuit homeostasis. Concurrently, genes associated with innate immunity, macrophage activation, and pro-inflammatory signaling were upregulated, suggesting that C-LTMRs normally suppress neuroinflammatory cascades. Following paclitaxel treatment, control mice exhibited transcriptional changes consistent with adaptive, pro-resolving responses. In contrast, *C-LTMRs-DTR* mice failed to engage many of these recovery-associated pathways and instead displayed enhanced expression of genes linked to neutrophil activation and tissue stress responses. These alterations point to a failure in molecular reprogramming necessary for pain resolution, providing a mechanistic basis for the persistent hypersensitivity observed behaviorally.

Taken together, our findings refine the gate theory of pain by identifying C-LTMRs as specialized sensory gatekeepers that regulate not only acute inhibition of nociceptive signals, but also the transcriptional and cellular environments that determine long-term recovery. C-LTMRs suppress central sensitization, restrain neuroimmune activation, and promote transcriptional programs required for tissue repair and sensory normalization. Their absence leads to unopposed sensitization, impaired resolution, and a shift toward chronic pain. These results argue against the view that all low-threshold mechanoreceptors contribute equally to analgesia, and instead highlight a unique, non-redundant role for C-LTMRs in maintaining sensory balance. Therapeutically, these findings suggest that enhancing C-LTMR function, through behavioral, pharmacological, or neuromodulatory interventions, could offer a novel strategy to prevent or reverse maladaptive pain states following injury or chemotherapy.

## EXPERIMENTAL PROCEDURES

### Mice

Mice were maintained under standard housing conditions (22°C, 40% humidity, 12 hr light cycles, and free access to food and water). Mice at 8 to 12 weeks of age and of both sexes were used for experiments. Particular efforts were made to minimize the number of mice used in this study, as well as the stress and suffering to which they were subjected. All experiments were conducted in line with the European guidelines for the care and use of laboratory animals (Council Directive 86/609/EEC). All experimental procedures were approved by an independent ethics committee for animal experimentation (APAFIS), as required by the French law and in accordance with the relevant institutional regulations of French legislation on animal experimentation, under license number APAFIS #34501. All experiments were performed in accordance with the ARRIVE guidelines.

### Generation of mouse lines

Na_v_1.8^Ires−FLPo^ mice (generated by our lab, Malapert et al., 2024) were crossed with TH^2ACreERT^^2^ mice (Abraira et al., 2017) and Tau^ds-DTR^ (gift of Dr Martyn Goulding) to generate Na_v_1.8^Ires−FLPo/+^::TH^2ACreERT^^2/+^::Tau^ds-DTR/+^ named *C-LTMRs-DTR*mice. Na_v_1.8^Ires−FLPo/+^::Tau^ds-DTR/+^ were used as control.

Na_v_1.8^Ires−FLPo^ mice (generated by our lab, Malapert et al., 2024) were crossed with RC::FL-hM3Dq mice (gift of Dr Patricia Jensen, National Institute of Environmental Health Sciences; cf. Sciolino et al., 2016) to generate Nav1.8^Ires-FLPo^::RC::FL-hM3Dq (related to Fig. S2).

### Tamoxifen treatmeant

Tamoxifen (Sigma #T5648) was freshly prepared and dissolved in corn oil (Sigma #C8267) to a concentration of 20 mg/mL. Tamoxifen solution was administered by using oral gavage (Instech Laboratories™ Feeding tube, rodent oral gavage, stainless steel, 22ga x 25mm, straight, sterile_Brand: Instech Laboratories™ FTSS-22S-25), once daily for five consecutive days, to 3-week-old mice. Both *C-LTMRs-DTR* and control mice were treated.

### Diphteria toxin treatmeant

Diphtheria Toxin (Sigma #322326-1MG) was dissolved in water then aliquoted and stored at −80°C. Diphteria Toxin solution was freshly prepared and administrated by using I.P (50µg/kg) on 2 days; separated by 48 h. Behavioral tests were performed 7 to 10 weeks after the initial DT injection. Both *C-LTMRs-DTR* and control mice were treated.

### Tissue processing for immunofluorescence (IF) and *in situ* hybridization (ISH)

Mice of both sexes were used for all experiments. Mice were deeply anesthetized with 100 mg/kg ketamine plus 10 mg/kg xylazine, and were intracardially perfused with an ice-cold solution of phosphate-buffered saline (PBS) followed by 30 ml ice-cold 4% paraformaldehyde in PBS. Their tissues were dissected and post-fixed by overnight incubation in the same fixative at 4°C.

DRG, JNG, ceacliac ganglion, and brain tissues were transferred to 30% (w/v) sucrose in PBS for cryoprotection and incubated at 4°C until they sank. They were then frozen in OCT medium and stored at −80 °C. Samples with a thickness of 12 μm (DRG, JNG) or 20µm (Cealiac), were cut with a standard cryostat (Leica). All these tissue sections were mounted on Superfrost slides and kept at −80°C until their use for IHC experiments.

Brain and the lumbar segment of the spinal cord were mounted in a small 3% agarose block. Sections with a thickness of 80 μm were cut with a Leica VT1200S vibratome. These sections were collected in a six-well plate filled with PBS and stored at 4°C until their use for IHC experiments.

### Immunofluorescence

For immunostaining, sections were incubated for 1 h at room temperature in PBS-10% (vol/vol) donkey serum (Sigma), 3% (weight/vol) bovine albumin (Sigma), 0.4% Triton X-100 and then overnight at 4°C with primary antibodies diluted in the same blocking solution. The primary antibodies used in this study were chicken anti-GFP (1:1000, Thermo Fisher Scientific, A10262); rabbit anti-P2X3 (1:1000 Neuromics Cat# RA10109, RRID:AB_2157931); rabbit anti-CGRP (1:1000, Cabiochem, PC205L, for DRG staining); rabbit anti-neurofilament M (145 kDa) (1:1000, Sigma-Aldrich AB1987); rabbit anti-TH (1:500, Sigma-Aldrich AB152); rat anti-TAFA4 (1:2000, a gift from Sophie Ugolini (CIML)); goat anti-TRKC (1:500, R and D Systems Cat# AF1404, RRID:AB_2155412); guinea pig anti-VGLUT3 (1:1000, Synaptic Systems, 135204). After three washes for 5 minutes each in 1xPBS, sections were incubated for 1 h at room temperature with secondary antibodies diluted in the blocking solution described above. The corresponding donkey anti-chicken, anti-rat, anti-rabbit, anti-goat or anti-guineapig Alexa 488-, 555-, or 647-conjugated secondary antibodies (1:500, Thermo Fisher Scientific) were used for the detection of primary antibody binding. Isolectin B4 conjugates with AlexaFluorR 647 dye were used at a dilution of 1:200 (Thermo Fisher Scientific I32450).

Tissues were washed (3 times in 1xPBS) and mounted in ImmuMount Reagent. Images were acquired with an AxioImager M2 (Zeiss) fluorescence microscope with a 20x/0,8 objective and contrast was adjusted with Fiji software.

#### Cell count

The entire L3 DRG was sectioned at 12 µm thickness using step serial sectioning and distributed across seven slides, ensuring that each slide contained a representative cross-section of the entire ganglion. Quantification of TH-, CGRP-, or P2X3-positive neurons was performed on all DRG sections present on a single slide from the L3 level. To estimate the total number of TH^+^, CGRP^+^, or P2X3^+^ neurons in a whole L3 DRG, the count obtained from that single slide was multiplied by seven. All analyses were conducted blind to the genotype of the animals.

### *In situ* hybridization

RNA probes were synthesized with gene-specific PCR primers and cDNA templates from mouse DRG. *In situ* hybridization was performed with digoxigenin-labeled probes (Roche, cat# 11277073910). Probes were incubated with the slides overnight at 55°C and the slides were then incubated with the horseradish peroxidase-conjugated anti-digoxigenin antibody 1:500 (Roche, Cat#11207733910; RRID:AB_514500). Final detection was achieved with TSA-Cy3 at a dilution of 1:50 (Perkin Elmer Life Sciences, FP1170). The following oligonucleotides were used for the nested PCRs for probe synthesis:

Ceacam10 F1 gactactgctcacagcctcact
Ceacam10 R1 cctactgctttttagcgtgaac
Ceacam10 F2 tggtacaagggaaacagtgg
Ceacam10 R2 TAATACGACTCACTATAGGGggcattagggtatgatcgaagt

### Electron Microscopy (EM)

#### Tissue preparation for ultrastructural morphology

Mice of both sexes were used for all experiments. Mice were deeply anesthetized with 100 mg/kg ketamine plus 10 mg/kg xylazine and perfused with Ringer solution followed by 1% PAF + 2% glutaraldehyde in 0.1 M phosphate buffer, pH 7.4. Lumbar spinal cord segments were dissected out and postfixed for 2 h in the same aldehyde mixture. Coronal sections were cut on a vibratome (Leica VT1000S) at a thickness of 80 μm.

#### EM embedding

Spinal cord sections were post-fixed in osmium ferrocyanide for 1 h at 4°C, dehydrated in graded acetone, incubated in acetone/Spurr resin (1:1; 30 min), acetone/Spurr resin (1:2; 30 min) and Spurr resin overnight at room temperature. Finally, sections were flat-embedded in Spurr resin (24 h, at 70°C). Ultrathin sections were cut with an ultramicrotome (EM UC6, Leica) and collected on uncoated nickel grids (200 mesh).

#### EM post-embedding immunostaining

Ultrathin sections were doubled immunostained following a conventional post-embedding protocol ([Salio et al., 2005](https://www-cell-com.insb.bib.cnrs.fr/cell-reports/fulltext/S2211-1247(19)31698-5?_returnURL=https%3A%2F%2Flinkinghub.elsevier.com%2Fretrieve%2Fpii%2FS2211124719316985%3Fshowall%3Dtrue)). Sections were treated for 2 min with a saturated aqueous solution of sodium metaperiodate (Sigma), rinsed in 1% Triton X-100 in Tris buffered saline (TBS) 0.5 M, and incubated for 1 h in 10% normal serum. Grids were then incubated overnight on drops of a mixture of primary antibodies at optimal dilutions. After rinsing in TBS, they were incubated in a mixture of the appropriate secondary antibodies gold conjugates (1:15; BBI Solutions, Cardiff, United Kingdom). They were then transferred into drops of 2.5% glutaraldehyde in cacodylate buffer 0.05 M pH 7.4 for 10 min, and finally washed in distilled water. Sections were further counterstained for 30’’ with Uranyl Less EM Stain and for 30’’ with Lead citrate (Electron Microscopy Sciences, Hatfield, PA, USA).

Sections were observed with a JEM-1010 transmission electron microscope (Jeol, Tokyo, Japan) equipped with a side-mounted CCD camera (Mega View III, Olympus Soft Imaging System, Brandeburg, Germany).

The following primary antibodies were used for EM immunostaining: rabbit anti-VGLUT3 (1:20, Abcam, Cat# ab23977 RRID: AB_2270290), IB4-biotin conjugate (1:20 Sigma-Aldrich Cat# L2140, RRID:AB_2313663.

Immunostained sections were observed with a JEM-1010 transmission electron microscope (Jeol, Tokyo, Japan) equipped with a side-mounted CCD camera (Mega View III, Olympus Soft Imaging System, Brandeburg, Germany).

#### EM Quantification of Glomeruli

We counted the number of glomeruli in lamina II to assess whether there was a loss after DT-selective ablation of C-LTMRs. We focused our analysis on counting GIa from control vs *C-LTMRs-DTR* animals in (i) plain ultrathin sections; and (ii) sections immunolabeled for IB4 and VGLUT3. Ten sections/animal for control and *C-LTMRs-DTR* mice were examined in studies that were carried out by an operator unaware of the experimental group. More precisely, the experimenter directly counted the number of glomeruli within all the 90 × 90 µm squares of 200 mesh EM grids that were occupied by relevant tissue. Quantitative analysis was performed using the ImageJ software (NIH, Bethesda, USA) and Graph Pad Prism 6 (GraphPad Software, San Diego, CA, USA).

### Pain models

#### Paw incision

Paw incision surgery was performed as described by Brennan (Brennan et al., 1999). Mice were anesthetized with ketamine (40 mg/kg IP) and xylazine (5 mg/kg IP) and a longitudinal incision was made through the skin and fascia of the right hind paw. Forceps were used to elevate the flexor digitorum brevis muscle longitudinally and an incision was made through the muscle with a scalpel, to cut it into two halves. The wound was closed with sutures, and the animals were allowed to recover and returned to their cages.

#### Paclitaxel treatment

Paclitaxel (Sigma 580555-5MG) was dissolved in a mixture of 1:1 [1 volume ethanol/1 volume Kolliphor-620 (Kolliphor-EL chez Sigma C5135). Paclitaxel solution was extemporaly prepared at a concentration of 0.4 mg/mL by diluting the 5 mg/ml stock solution with 0.9% NaCl. The mice received an intraperitoneal injection of paclitaxel (4 mg/kg) every two days for a total of four injections. The last injection was administered 72 hours after the third injection. Both *C-LTMRs-DTR* and control mice were treated.

### Behavior

All behavioral assays were conducted on 11- to 14-week-old mice. Animals were acclimated to their testing environment for 45-60 minutes before each experiment, and all experiments were performed at room temperature (~22°C). Experimenters were blind to the treatments used. Mice of both sexes were used for all experiments.

#### Open-field test

The open-field apparatus consists of an empty square arena (40×40×35 cm), surrounded by walls to prevent animal from escaping. Light inside the arena was uniform and kept at approximately 100 lux throughout the tests. Control and cre positive mice were individually placed in the center of the arena and their behavior was recorded using the EthoVision XT16 video-tracking system (Noldus) over a 10-minute period. The time spent grooming and rearing, the total distance traveled, and the total amount of time spent in the peripheral borders and in the center were recorded.

#### Rotarod test

A rotarod apparatus (LSI Letica Scientific Instruments) was used to explore coordinated locomotor and balance function in mice. Mice were placed on a rod that slowly accelerated from 4 rpm to 44 rpm over 5 minutes and the latency to fall off during this period was recorded.

The test was conducted over 3 consecutive days. Each day, the animals were tested 3 times separated by at least a 5-minute resting period.

#### Thermal gradient test

Response to temperature Gradient assay were performed as described in (Moqrich et al., 2005) but using Bioseb apparatus.

#### Tape test

Mice are allowed to acclimate in a circular plexiglass container for 5 minutes. A 3 cm piece of common lab tape was then applied gently to the back of the mouse such that it sticks to the mouse. Mice are then observed for 5 minutes. The latency before the first response is recorded and the total number of responses to the tape are counted. A response is scored when the mouse stops moving and bites or scratches the piece of tape or shows a visible “wet dog shake” motion in an attempt to remove the foreign object on its back.

#### Oil droplet test

This test was performed as described in (Zhang et al., 2024). Briefly, oil droplet stimuli, 16-18 µl of sunflower seed oil (Sigma #S5007) were applied to the neck of the mice using a glass Pasteur pipette. A response is scored when the mouse shows a visible “wet dog shake” motion in an attempt to remove the oil on its neck.

#### Von Frey’s test

Mice were placed in plastic chambers on a wire mesh grid and stimulated with von Frey filaments (Bioseb) by the up-down method (Chaplan et al., 1994) starting with a 1g filament, and using 0.07 and 2g filaments as the cutoffs.

### Dry ice test

The mice are acclimated on a glass plate (8 mm thick float borosilicate Pyrex) in transparent plastic enclosures separated by opaque black partitions for 30 minutes to one hour. Resting mice, but not sleeping mice, are tested by placing a dry ice pellet under the hindpaw on the glass plate. The dry ice should be placed in the center of the hind paw, being careful to avoid the distal joints and ensuring good contact between the paw and the glass. Withdrawal latency is measured with a stopwatch and defined as any action aimed at moving the paw away from the cold glass, either vertically or horizontally. There is an interval of at least 15 minutes between tests on the same paw. These intervals were chosen empirically to allow sufficient time for the mouse to return to a resting state after stimulation. Each paw is measured at least three times. The maximum time allowed for withdrawal is 20 seconds to avoid potential tissue damage. Trials in which the animal does not withdraw within 20 seconds are repeated. During the second test, if there is no withdrawal within the threshold, the value is recorded as 20 seconds.

### High-throughput RNA sequencing and analyses

Control and *C-LTMRs-DTR* DRG or DHSC RNAs from naive and 40 days post paclitaxel mice (only males), were extracted in experimental quadruplate from individual mice. High quality RNA (RIN > 8) was used for sequencing. RNA quality was measured using Agilent RNA 6000 Pico Kit. RNA-seq libraries were prepared using Watchmaker mRNA Library Prep Kit (Watchmaker Genomics) in order to produce paired end reads of 100 pb. All libraries were validated for concentration and fragment size using Agilent DNA1000 chips. Sequencing was performed on a NovaSeqX (Illumina) and quality control performed using FastQC (https://www.bioinformatics.bbsrc.ac.uk/projects/fastqc). Sequences were uniquely mapped to the mm39 genome using STAR (Dobin et al., 2012) (version 2.7.11b) using default values and paired-end mode. Reads mapping to gene exons (GRCm39 GCF_000001635.27 NCBI RefSeq assembly) were counted using featureCounts (Liao et al., 2014) (C version 1.4.6-p2). Differential gene expression was performed using exon counts from biological replicates using the EdgeR BioConductor R package (Robinson et al., 2010) (version 4.2.2), using a 5 % false discovery rate (FDR) cutoff. Heat-maps were generated using Heatmapper on-line software (https://heatmapper.ca). Venny diagrams were generated using Venny on-line software (https://bioinfogp.cnb.csic.es/tools/venny/). Functional analysis was performed using Metascape software (Zhou et al., 2019).

### Software

Some figures elements were generated using BioRender on-line software (https://www.biorender.com/)

### Quantification and statistical analysis

Results were expressed as mean +/− SEM. Quantitative and statistical analyses were performed by using the GraphPad Prism 7 (GraphPad Software, La Jolla, CA) and were indicated in each figure. The Shapiro–Wilk test was used to assess the normality of the data. Statistical significance was set to *P, 0.05, **P, 0.01,***P, 0.001, and ****P, 0.0001.

## AUTHORS’ CONTRIBUTION

G.R characterized the mouse model, managed the moue colony and generated most of the data presented in the manuscript. He also generated all the figures and managed the writing of the materials and methods section, K.M sat up the CIPN model and generated the related data, contributed to the immunostaining and RNA-seq data. A.C generated the paw incision data. P.M generated the Nav1.8-FLP mouse model. C.S performed the electron microscopy experiments and generated data that will be published elsewhere. A.S analysed the raw RNA-seq data, A.R managed the progression of the project, contributed to the generation and analysis of the the RNA-seq data. A.M designed the project and wrote the manuscript. All authors contributed to editing the manuscript.

## ACKNOWLEGMENTS

We are grateful to the members of the Moqrich lab at IBDM for the scientific discussions. The IBDM imaging and animal facilities for assistance. This work was funded by the ANR Sensorimmune and by institutional funding from the CNRS and Aix-Marseille-Université to IBDM.

