## Supplementary material for "C-LTMRs Regulate Thermosensation and Gate the Transition from Acute to Chronic Pain": Suplemental figures and legends

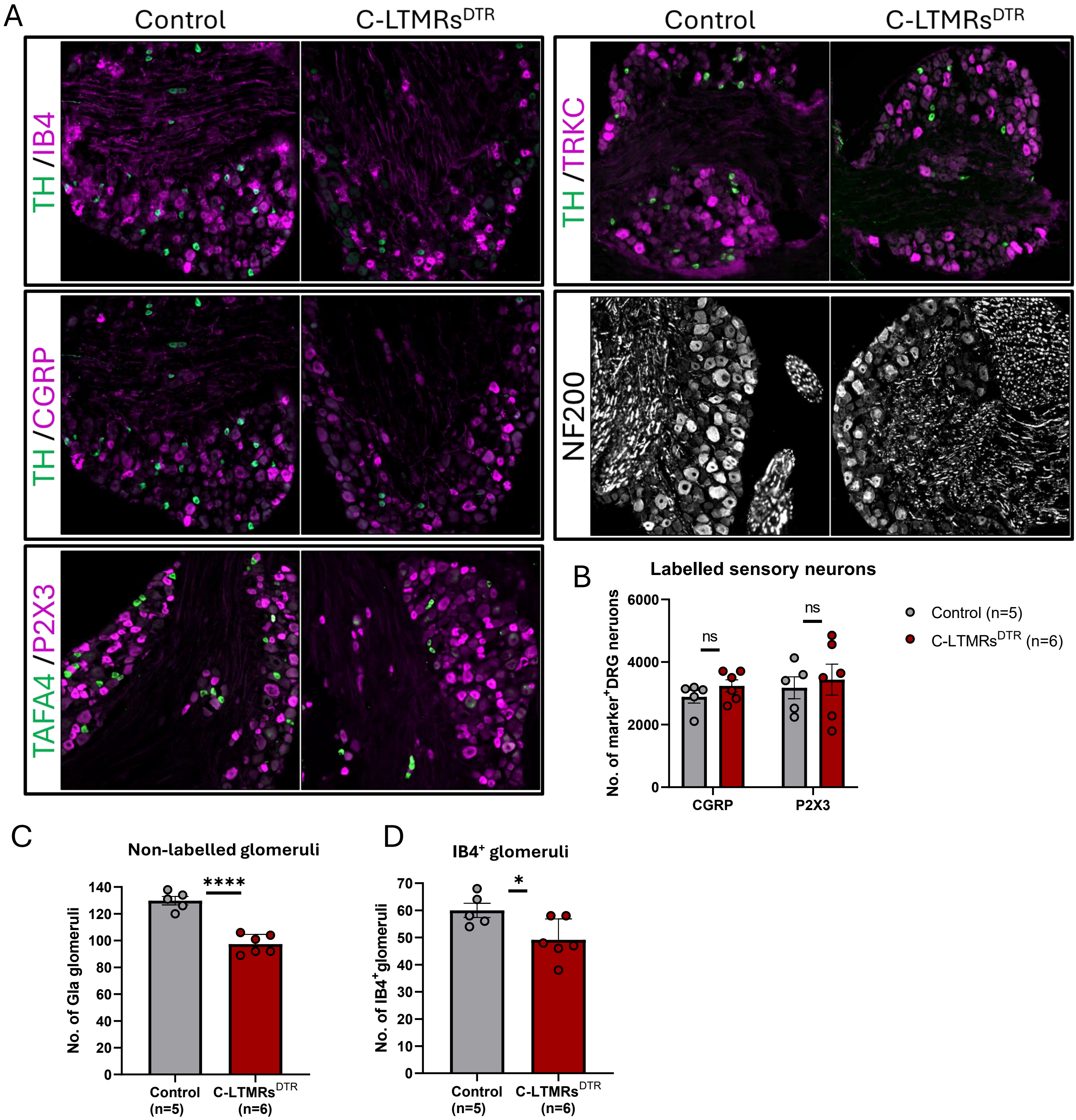
A
Control
C-LTMRs$^{DTR}$
Control
C-LTMRs$^{DTR}$
TH /IB4
TH /TRKC
TH /CGRP
NF200
TAFA4 /P2X3
B
Labelled sensory neurons
No. of marker$^{+}$DRG neruons
6000
4000
2000
0
ns
ns
CGRP
P2X3
Control (n=5)
C-LTMRs$^{DTR}$ (n=6)
C
Non-labelled glomeruli
No. of Gla glomeruli
140
120
100
80
60
40
20
0
****
Control (n=5)
C-LTMRs$^{DTR}$ (n=6)
D
IB4$^{+}$ glomeruli
No. of IB4$^{+}$glomeruli
70
60
50
40
30
20
10
0
*
Control (n=5)
C-LTMRs$^{DTR}$ (n=6)

**Figure S1. Selectivity of the C-LTMRs deletion in the DRG and the DHSC**

**(A)** Representative images of lumbar (L3) DRG sections stained with cell type markers in control and *C-LTMRs-DTR* mice.

**(B)** Quantification of the total number of lumbar (L3) $CGRP^+$ DRG neurons (t-test) and $P2X3^+$ DRG neurons (Mann Witney test) in control (n=5) and *C-LTMRs-DTR* mice (*n*=6). Data are presented as mean ± SEM. Dots represent individual animals.

**(C-D)** Quantification of (C) non-labelled GIa glomeruli and (D) $IB4^+$ glomeruli in the DHSC, in control (n=5) and *C-LTMRs-DTR* mice (n=6). Data are presented as mean ± SEM. Dots represent individual animals. Statistical comparisons were performed using T-Test, *$P < 0.05$ *et* ****$P < 0.0001$.

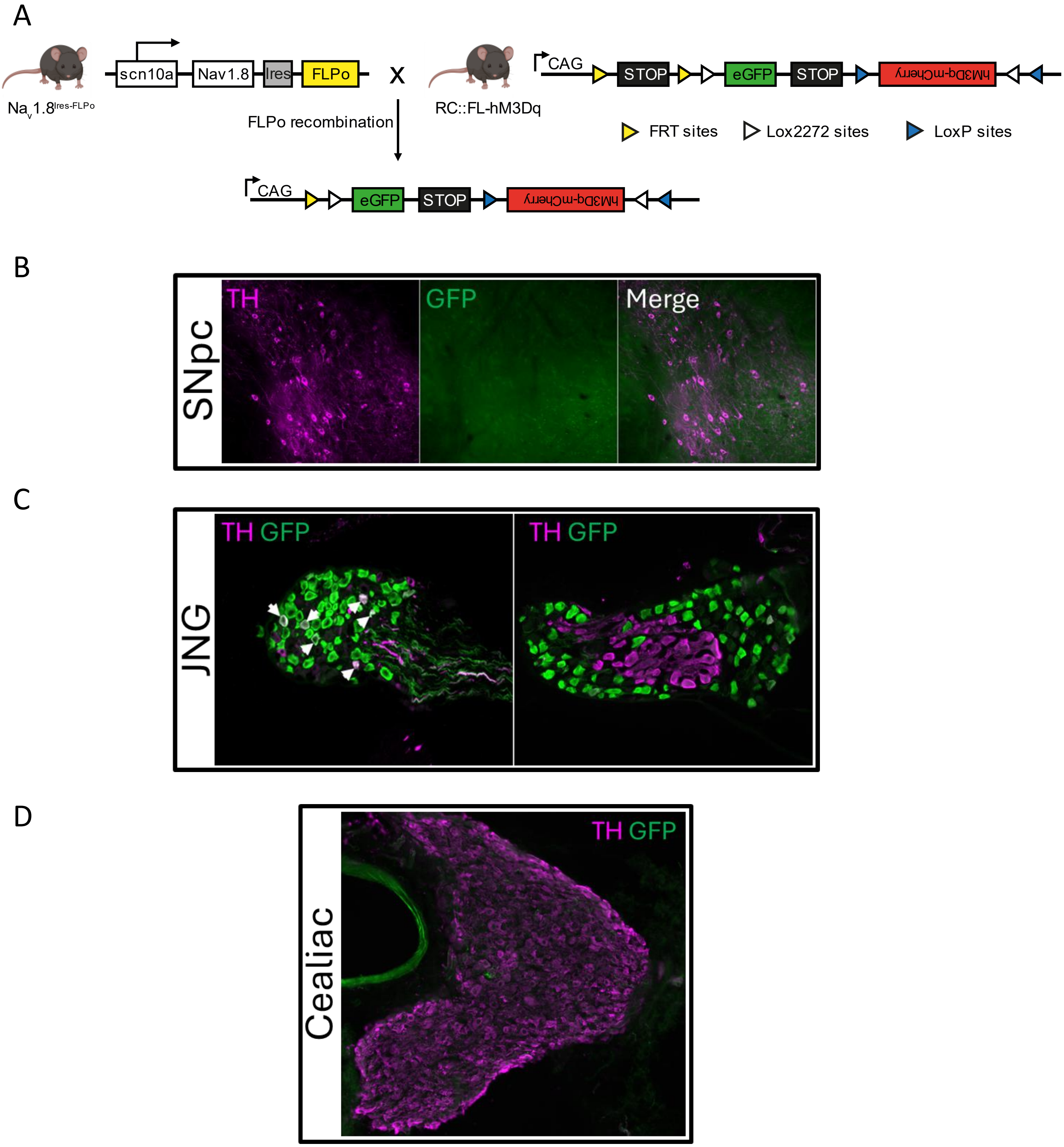
A
$Na_v1.8^{Ires-FLPo}$
scn10a
Nav1.8
Ires
FLPo
x
RC::FL-hM3Dq
CAG
STOP
eGFP
STOP
hM3Dq-mCherry
FLPo recombination
FRT sites
Lox2272 sites
LoxP sites
CAG
eGFP
STOP
hM3Dq-mCherry
B
SNpc
TH
GFP
Merge
C
JNG
TH GFP
TH GFP
D
Cealiac
TH GFP

**Figure S2. Selectivity of the Nav1.8-TH signature to C-LTMRs**

**(A)** Schematic representation of the strategy for mating between a Nav1.8$^{Ires\text{-}FLPo}$ mouse and a reporter RC::FL-hM3Dq mouse. In the progeny of this cross, the FLPo will excise the FRT-flanked transcription STOP cassette, allowing the expression of EGFP in the Nav1.8 lineage cells.

**(B–D)** TH (magenta) and GFP (green) immunostaining on (B) sagittal brain section at the level of the Substantia Nigra Pars Compacta (SNpc), (C) jugular-nodose-ganglia (JNG) section and **(D)** Cealiac ganglion section from a $Na_v1.8^{Ires\text{-}FLPo}$::RC-FL-hM3Dq mouse.

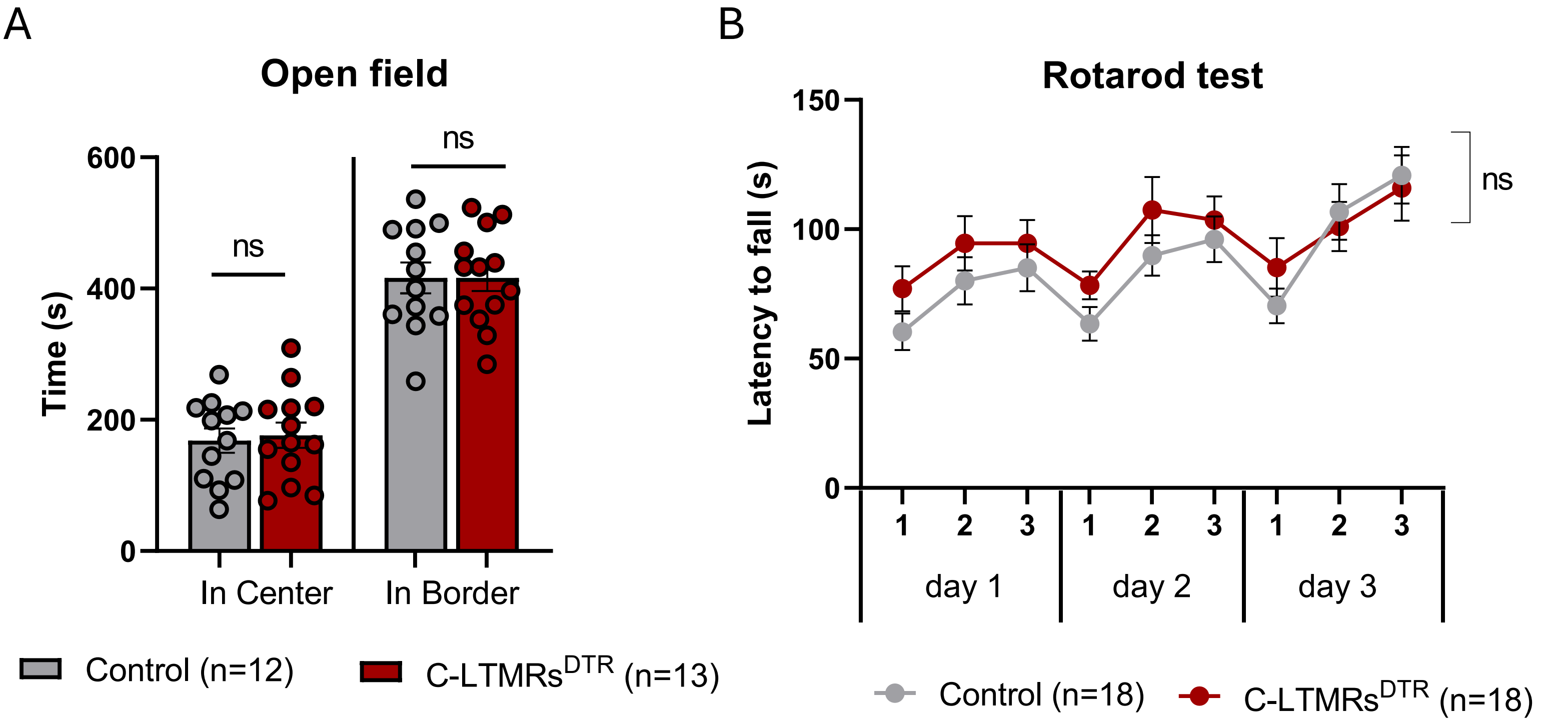
A
Open field
ns
ns
600
400
200
0
Time (s)
In Center
In Border
Control (n=12)
C-LTMRs^DTR (n=13)
B
Rotarod test
150
100
50
0
Latency to fall (s)
ns
1
2
3
1
2
3
1
2
3
day 1
day 2
day 3
Control (n=18)
C-LTMRs^DTR (n=18)

**Figure S3. Partial ablation C-LTMRs ablation does not induce anxiety behavior or disturbances in sensorimotor coordination**

(A-B) *C-LTMRs-DTR* mice display unaltered phenotype compared to control mice in (A) anxiety with openfield test (n=12 control, n=13 *C-LTMRs-DTR*) (B) in motor coordination with rotaroad test (n=18 in each group)

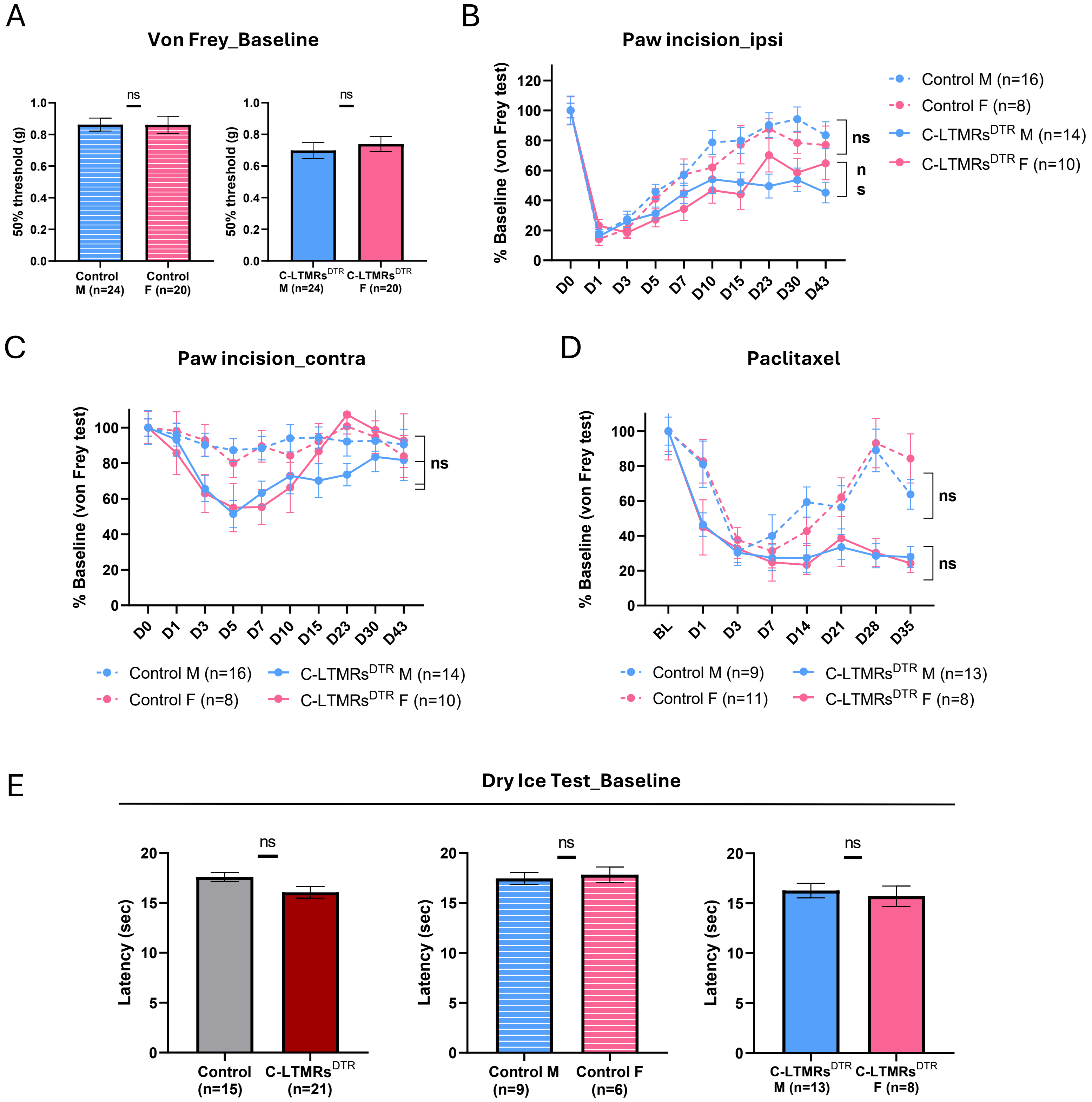
A
Von Frey_Baseline
50% threshold (g)
ns
Control M (n=24)
Control F (n=20)
C-LTMRs$^{DTR}$ M (n=24)
C-LTMRs$^{DTR}$ F (n=20)
B
Paw incision_ipsi
% Baseline (von Frey test)
D0 D1 D3 D5 D7 D10 D15 D23 D30 D43
Control M (n=16)
Control F (n=8)
C-LTMRs$^{DTR}$ M (n=14)
C-LTMRs$^{DTR}$ F (n=10)
C
Paw incision_contra
D
Paclitaxel
BL D1 D3 D7 D14 D21 D28 D35
Control M (n=9)
Control F (n=11)
C-LTMRs$^{DTR}$ M (n=13)
C-LTMRs$^{DTR}$ F (n=8)
E
Dry Ice Test_Baseline
Latency (sec)
Control (n=15)
C-LTMRs$^{DTR}$ (n=21)
Control M (n=9)
Control F (n=6)
C-LTMRs$^{DTR}$ M (n=13)
C-LTMRs$^{DTR}$ F (n=8)

**Figure S4. Males and Females of both genotypes display no differences for mechanical and cold plantar sensitivity in naive conditions and following injury**

**(A)** Baseline mechanical thresholds measured by Von Frey testing show no differences between males (n=24) and females (n=20) of both genotypes (unpaired t-test).

**(B-D)** Males and females of both genotypes behave in a similar manner across all time point in terms of mechanical sensitivity of (B) the ipsi or C) the contra lateral side before and following paw incision and D) before and following paclitaxel. Results are expressed as mean ± SEM for indicated mice number (two-way repeated measures ANOVA).

**(E)** No differences were observed between control (n=15) and *C-LTMRs-DTR* mice (n=21) in terms of cold sensitivity, at the baseline level. Males and females of both genotypes behave in a similar manner. Results are expressed as mean ± SEM for indicated mice number (unpaired t-test).

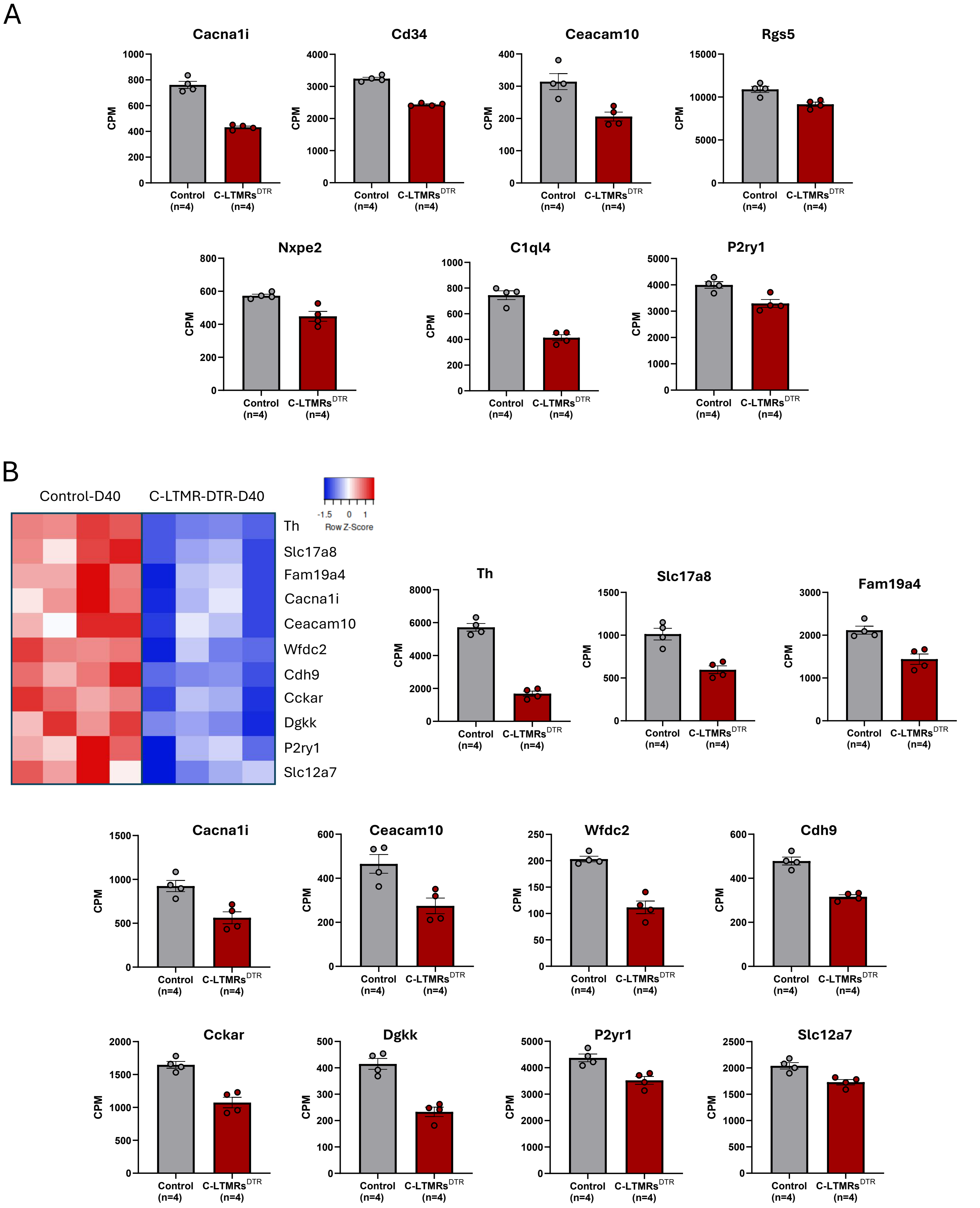
A
Cacna1i
Cd34
Ceacam10
Rgs5
Nxpe2
C1ql4
P2ry1
CPM
Control (n=4)
C-LTMRs^DTR (n=4)
B
Control-D40
C-LTMR-DTR-D40
Row Z-Score
-1.5 0 1
Th
Slc17a8
Fam19a4
Cacna1i
Ceacam10
Wfdc2
Cdh9
Cckar
Dgkk
P2ry1
Slc12a7
Th
Slc17a8
Fam19a4
Cacna1i
Ceacam10
Wfdc2
Cdh9
Cckar
Dgkk
P2yr1
Slc12a7

**Figure S5. *C-LTMRs-DTR* mice exhibit a drastic decrease in the expression of many C-LTMRs-specific and enriched genes**

**(A**) Counts Per Million (CPM) obtained after RNA-Seq for the C-LTMR-specific and enriched genes presented in Fig.4A, in control (gray) and *C-LTMRs-DTR* (red) mice. Results are expressed as mean ± SEM for indicated mice number. Dots represent individual animals.

**(B)** Heatmap representing the expression of indicated C-LTMRs-specific and enriched genes in Control and *C-LTMRs-DTR* mice, 40 days following paw incision; and graphical representations of the level of expression (CPM) of these genes. Results are expressed as mean ± SEM for indicated mice number. Dots represent individual animals.

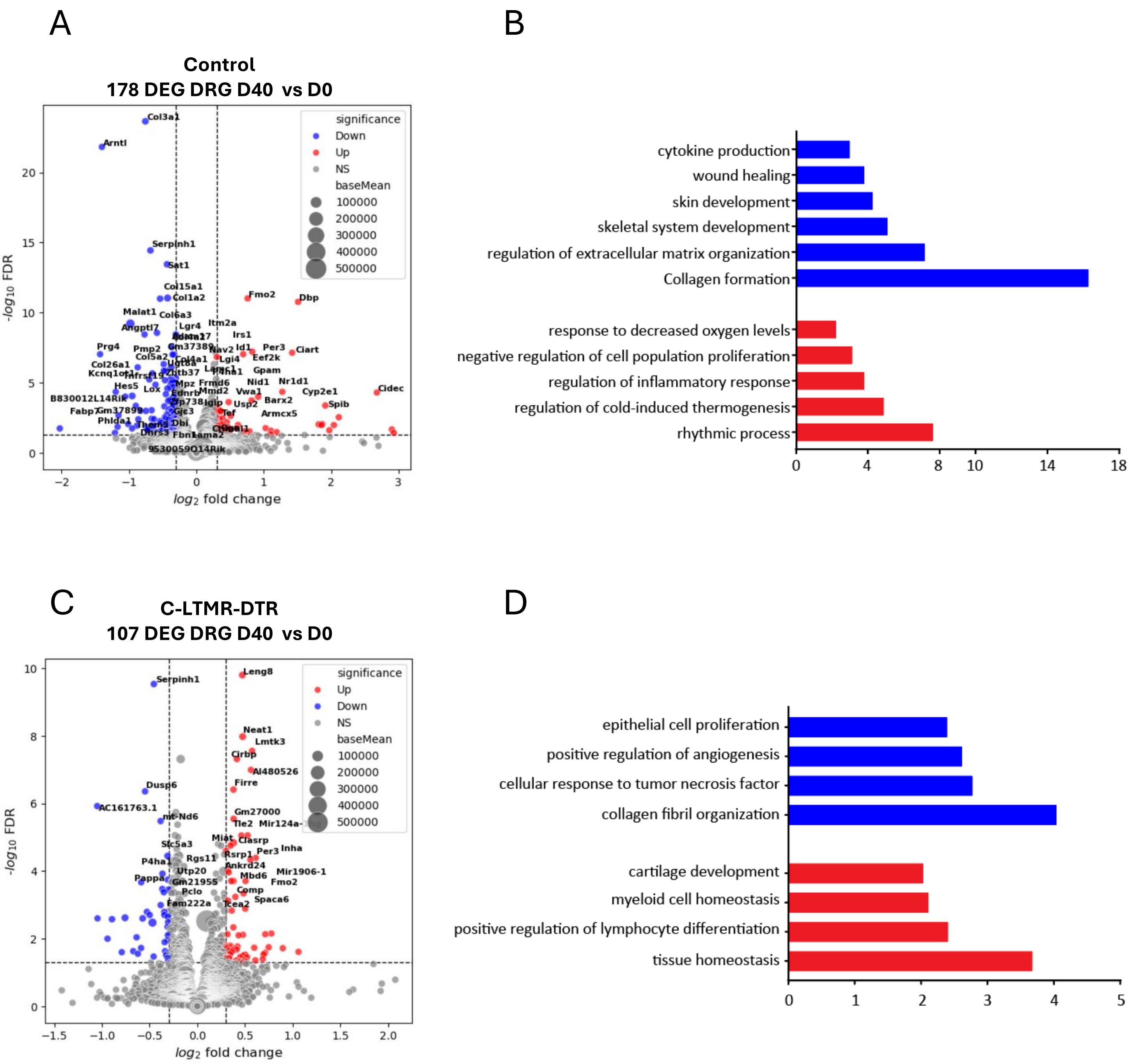

A
Control
178 DEG DRG D40 vs D0
significance
Down
Up
NS
baseMean
100000
200000
300000
400000
500000
$-log_{10}$ FDR
$log_2$ fold change
Col3a1
Arntl
Serpinh1
Sat1
Col15a1
Col1a2
Fmo2
Dbp
Malat1
Col6a3
Angptl7
Lgr4
Itm2a
Irs1
Prg4
Pmp2
Per3
Ciart
Col5a2
Col4a1
Id1
Eef2k
Col26a1
Kcnq1ot1
Lgi4
Gpam
Hes5
Lox
Mpz
Frmd6
Nid1
Nr1d1
Cidec
Ednrb
Mmd2
Vwa1
Cyp2e1
B830012L14Rik
Usp2
Barx2
Spib
Fabp7
Gm37899
Tef
Armcx5
Phlda1
Dbi
Gjc3
Fbn1
Lama2
9530059O14Rik
B
cytokine production
wound healing
skin development
skeletal system development
regulation of extracellular matrix organization
Collagen formation
response to decreased oxygen levels
negative regulation of cell population proliferation
regulation of inflammatory response
regulation of cold-induced thermogenesis
rhythmic process
C
C-LTMR-DTR
107 DEG DRG D40 vs D0
significance
Up
Down
NS
baseMean
100000
200000
300000
400000
500000
$-log_{10}$ FDR
$log_2$ fold change
Serpinh1
Leng8
Neat1
Lmtk3
Cirbp
AI480526
Dusp6
Firre
AC161763.1
mt-Nd6
Gm27000
Tle2
Slc5a3
Miat
Clasrp
Per3
Inha
P4ha1
Rgs11
Rsrp1
Utp20
Ankrd24
Mir1906-1
Pappa
Gm21955
Mbd6
Fmo2
Pclo
Comp
Spaca6
Fam222a
Tcea2
D
epithelial cell proliferation
positive regulation of angiogenesis
cellular response to tumor necrosis factor
collagen fibril organization
cartilage development
myeloid cell homeostasis
positive regulation of lymphocyte differentiation
tissue homeostasis

**Figure S6. Partial depletion of C-LTMRs modifies DRG response to paclitaxel**

A) Volacno plot representing up- and down-regulated DEG in the L3-L5 DRGs at day 40 (D40) after paclitaxel administration as compared to untreated (D0) Control mice (FDR5, Log2FoldChange > 0.3, n=4 samples per genotype). B) Up (red) and down (blue)-regulated biological processes and pathways in Control lumbar DRGs at D40 after paclitaxel administration, determined with Metascape software. C) Volacno plot representing up- and down-regulated DEG in the L3-L5 DRGs at day 40 (D40) after paclitaxel administration as compared to untreated (D0) *C-LTMRs-DTR* mice (FDR5, Log2FoldChange > 0.3, n=4 samples per genotype). D) Up (red) and down (blue)-regulated biological processes and pathways in *C-LTMRs-DTR* lumbar DRGs at D40 after paclitaxel administration, determined with Metascape software.

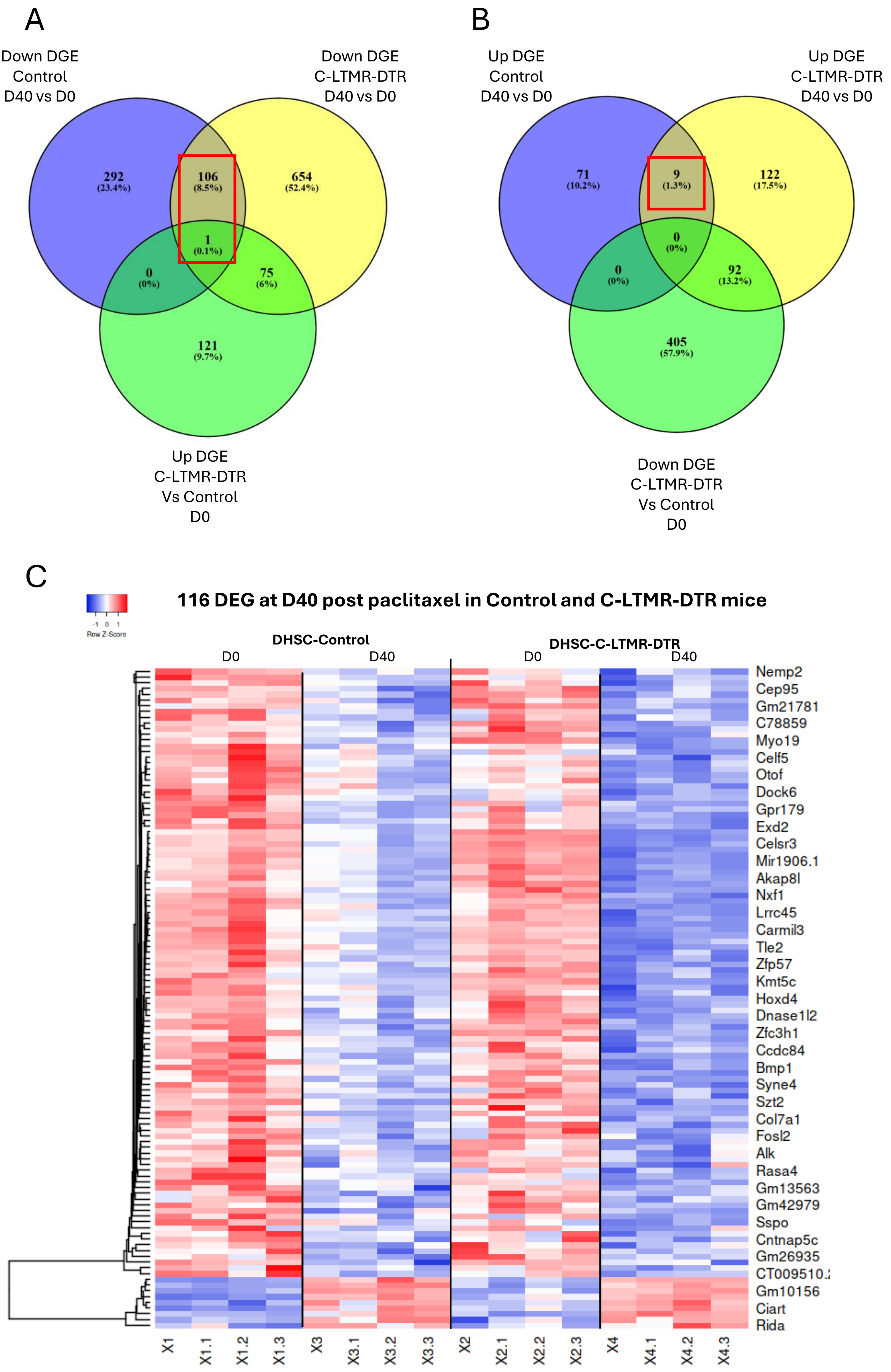
A
Down DGE
Control
D40 vs D0
Down DGE
C-LTMR-DTR
D40 vs D0
292
(23.4%)
106
(8.5%)
654
(52.4%)
1
(0.1%)
0
(0%)
75
(6%)
121
(9.7%)
Up DGE
C-LTMR-DTR
Vs Control
D0
B
Up DGE
Control
D40 vs D0
Up DGE
C-LTMR-DTR
D40 vs D0
71
(10.2%)
9
(1.3%)
122
(17.5%)
0
(0%)
0
(0%)
92
(13.2%)
405
(57.9%)
Down DGE
C-LTMR-DTR
Vs Control
D0
C
116 DEG at D40 post paclitaxel in Control and C-LTMR-DTR mice
-1 0 1
Row Z-Score
DHSC-Control
D0
D40
DHSC-C-LTMR-DTR
D0
D40
Nemp2
Cep95
Gm21781
C78859
Myo19
Celf5
Otof
Dock6
Gpr179
Exd2
Celsr3
Mir1906.1
Akap8l
Nxf1
Lrrc45
Carmil3
Tle2
Zfp57
Kmt5c
Hoxd4
Dnase1l2
Zfc3h1
Ccdc84
Bmp1
Syne4
Szt2
Col7a1
Fosl2
Alk
Rasa4
Gm13563
Gm42979
Sspo
Cntnap5c
Gm26935
CT009510.
Gm10156
Ciart
Rida
X1
X1.1
X1.2
X1.3
X3
X3.1
X3.2
X3.3
X2
X2.1
X2.2
X2.3
X4
X4.1
X4.2
X4.3

**Figure S7. Analysis of RNA-Seq data from DHSC to identify the sets of genes similarly affected by paclitaxel in Controls and *C-LTMRs-DTR* mice**

A-B) Venn diagrams were used to compare the sets of down and up DEG identified in the D40 vs D0 comparison in Control and *C-LTMRs-DTR* DHSC (FDR5, Lod2FoldChange >0.3, n=4 samples per experimental group). To identify the genes whose de-regulation was due to paclitaxel but not to the differences observed in naïve condition between *C-LTMRs-DTR* and control DHSC, the indicated sets of DEG at day 0 were included in the Venn diagram. Red rectangle shows the sets of genes (107) down and up (9)-regulated in both Control and *C-LTMRs-DTR* at day 40 after paclitaxel. C) Heatmap representing the expression levels of these 116 DEG which are common to both Control and *C-LTMRs-DTR* DHSC in each indicated experimental group.

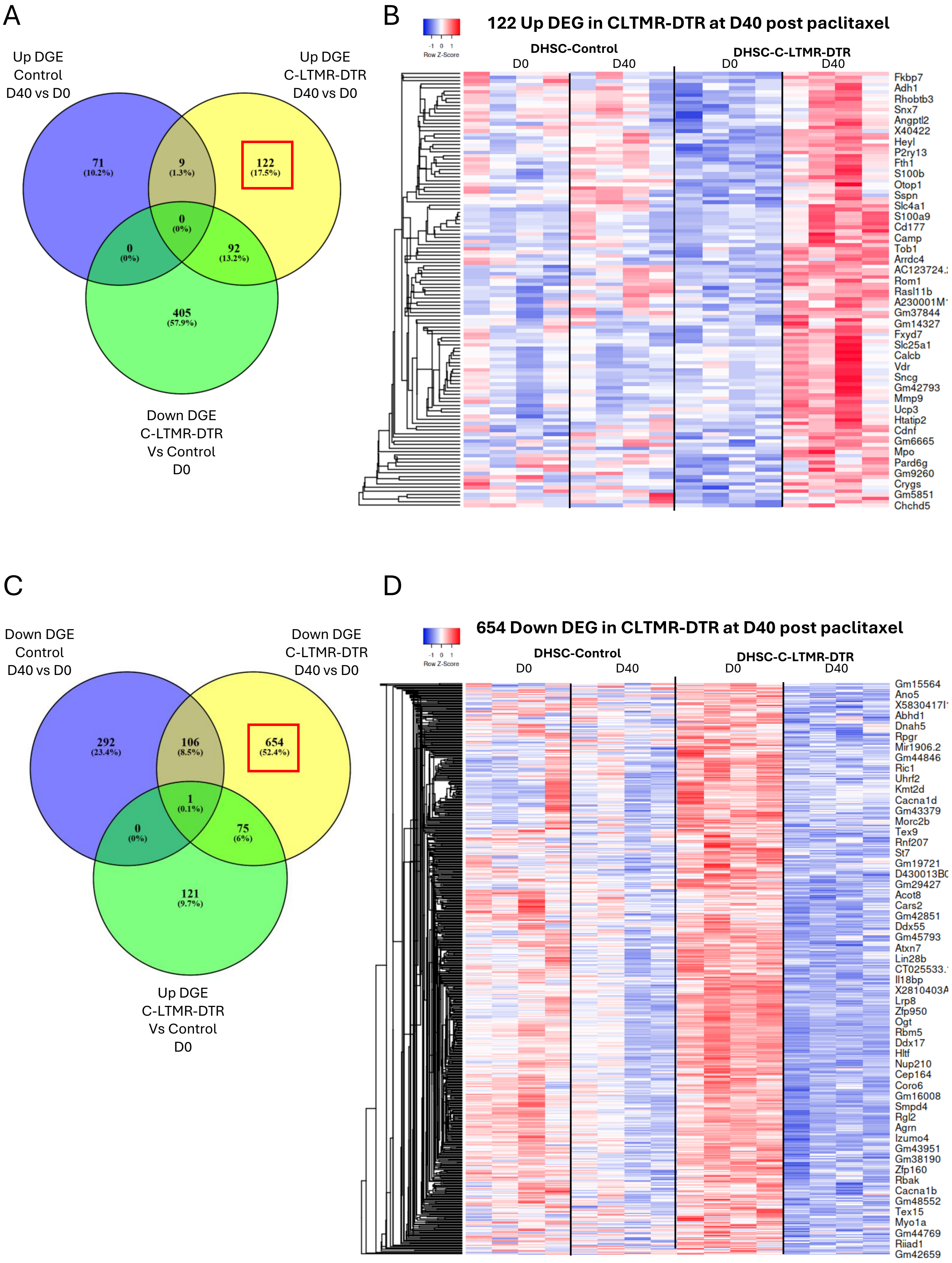

A
Up DGE Control D40 vs D0
Up DGE C-LTMR-DTR D40 vs D0
Down DGE C-LTMR-DTR Vs Control D0
71 (10.2%)
9 (1.3%)
122 (17.5%)
0 (0%)
0 (0%)
92 (13.2%)
405 (57.9%)
B
122 Up DEG in CLTMR-DTR at D40 post paclitaxel
-1 0 1
Row Z-Score
DHSC-Control
D0
D40
DHSC-C-LTMR-DTR
D0
D40
Fkbp7
Adh1
Rhobtb3
Snx7
Angptl2
X40422
Heyl
P2ry13
Fth1
S100b
Otop1
Sspn
Slc4a1
S100a9
Cd177
Camp
Tob1
Arrdc4
Rom1
Rasl11b
Gm37844
Gm14327
Fxyd7
Slc25a1
Calcb
Vdr
Sncg
Gm42793
Mmp9
Ucp3
Htatip2
Cdnf
Gm6665
Mpo
Pard6g
Gm9260
Crygs
Gm5851
Chchd5
C
Down DGE Control D40 vs D0
Down DGE C-LTMR-DTR D40 vs D0
Up DGE C-LTMR-DTR Vs Control D0
292 (23.4%)
106 (8.5%)
654 (52.4%)
1 (0.1%)
0 (0%)
75 (6%)
121 (9.7%)
D
654 Down DEG in CLTMR-DTR at D40 post paclitaxel
-1 0 1
Row Z-Score
DHSC-Control
D0
D40
DHSC-C-LTMR-DTR
D0
D40
Gm15564
Ano5
Abhd1
Dnah5
Rpgr
Mir1906.2
Gm44846
Ric1
Uhrf2
Kmt2d
Cacna1d
Gm43379
Morc2b
Tex9
Rnf207
St7
Gm19721
Gm29427
Acot8
Cars2
Gm42851
Ddx55
Gm45793
Atxn7
Lin28b
Il18bp
Lrp8
Zfp950
Ogt
Rbm5
Ddx17
Hltf
Nup210
Cep164
Coro6
Gm16008
Smpd4
Rgl2
Agrn
Izumo4
Gm43951
Gm38190
Zfp160
Rbak
Cacna1b
Gm48552
Tex15
Myo1a
Gm44769
Riiad1
Gm42659

**Figure S8. Analysis of RNA-Seq data from DHSC to identify the sets of genes uniquely disrupted in *C-LTMRs-DTR* mice**

A) Venn diagrams were used to compare the sets of down and up DEG identified in the D40 vs D0 comparison in Control and *C-LTMRs-DTR* DHSC (FDR5, Lod2FoldChange >0.3, n=4 samples per experimental group). 122 up-regulated genes in *C-LTMRs-DTR* DHSC at D40 after paclitaxel (D40 vs D0) were identified by excluding those that were common with Control up-DEG D40 vs D0 and those that were already down-regulated at day 0 in *C-LTMRs-DTR* as compared to Control mice. B) Heatmap projection of these 122 genes across experimental groups. C) 654 down-regulated genes in *C-LTMRs-DTR* DHSC at D40 after paclitaxel (D40 vs D0) were identified by excluding those that were in common with Control down-DEG D40 vs D0 and those that were already up-regulated at day 0 in *C-LTMRs-DTR* as compared to Control mice. D) Heatmap projection of these 654 genes across experimental groups.

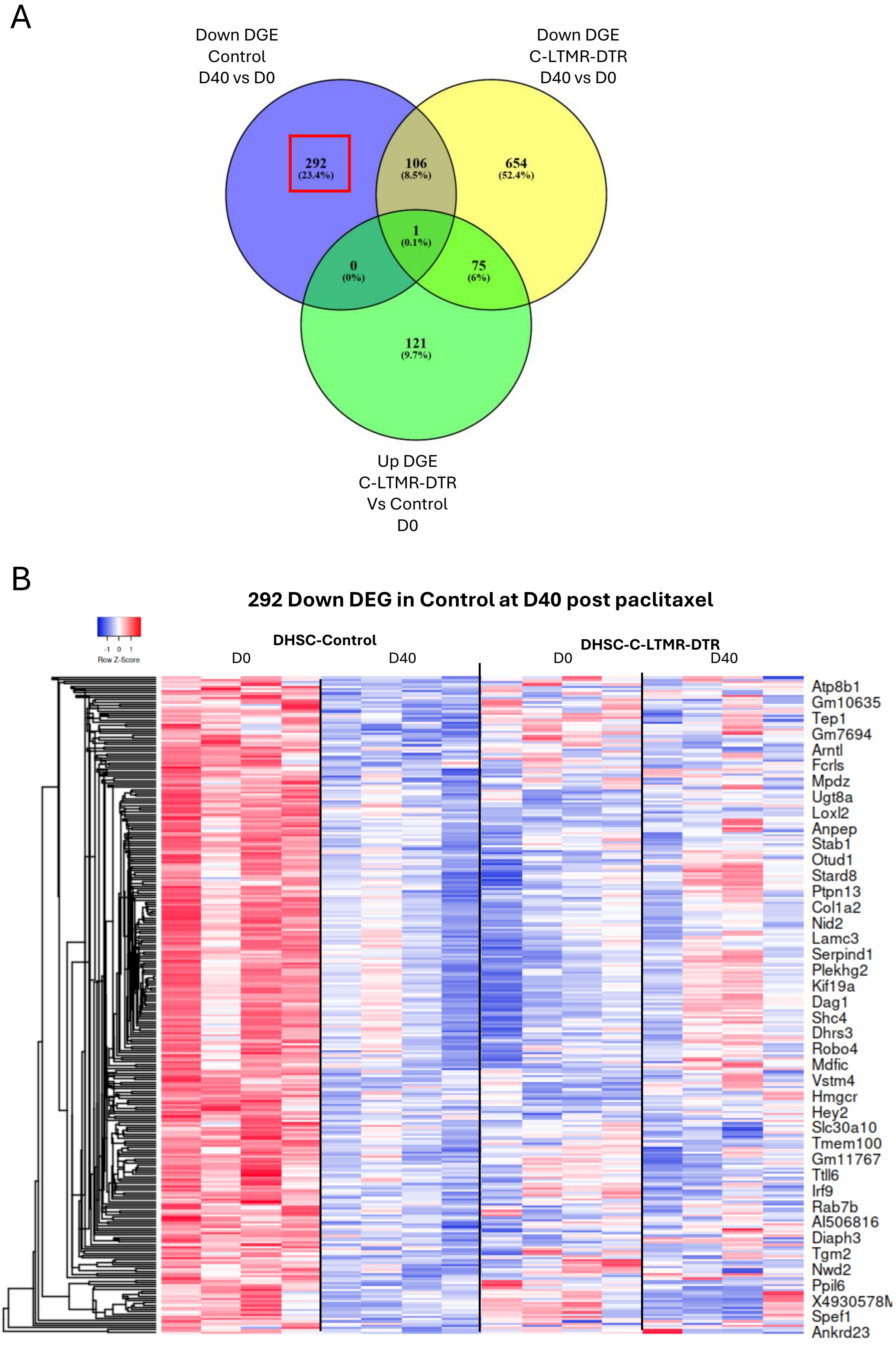
A
Down DGE
Control
D40 vs D0
Down DGE
C-LTMR-DTR
D40 vs D0
292
(23.4%)
106
(8.5%)
654
(52.4%)
1
(0.1%)
0
(0%)
75
(6%)
121
(9.7%)
Up DGE
C-LTMR-DTR
Vs Control
D0
B
292 Down DEG in Control at D40 post paclitaxel
Row Z-Score
-1 0 1
DHSC-Control
D0
D40
DHSC-C-LTMR-DTR
D0
D40
Atp8b1
Gm10635
Tep1
Gm7694
Arntl
Fcrls
Mpdz
Ugt8a
Loxl2
Anpep
Stab1
Otud1
Stard8
Ptpn13
Col1a2
Nid2
Lamc3
Serpind1
Plekhg2
Kif19a
Dag1
Shc4
Dhrs3
Robo4
Mdfic
Vstm4
Hmgcr
Hey2
Slc30a10
Tmem100
Gm11767
Ttll6
Irf9
Rab7b
AI506816
Diaph3
Tgm2
Nwd2
Ppil6
X4930578M
Spef1
Ankrd23

**Figure S9. Analysis of RNA-Seq data from DHSC to identify the sets of genes suppressed in Control mice in response to paclitaxel administration**

A) Venn diagrams were used to compare the sets of down and up DEG identified in the D40 vs D0 comparison in Control and *C-LTMRs-DTR* DHSC (FDR5, Lod2FoldChange >0.3, n=4 samples per experimental group). 292 genes were identified as being selective to the Control D40 vs D0 condition. B) Heatmap projection of these 292 genes across experimental groups.
